# Inferring viral transmission time from phylogenies for known transmission pairs

**DOI:** 10.1101/2023.09.12.557404

**Authors:** Emma E. Goldberg, Erik J. Lundgren, Ethan O. Romero-Severson, Thomas Leitner

## Abstract

When the time of an HIV transmission event is unknown, methods to identify it from virus genetic data can reveal the circumstances that enable transmission. We developed a single-parameter Markov model to infer transmission time from an HIV phylogeny constructed of multiple virus sequences from people in a transmission pair. Our method finds the statistical support for transmission occurring in different possible time slices. We compared our time-slice model results to previously-described methods: a tree-based logical transmission interval, a simple parsimony-like rules-based method, and a more complex coalescent model. Across simulations with multiple transmitted lineages, different transmission times relative to the source’s infection, and different sampling times relative to transmission, we found that overall our time-slice model provided accurate and narrower estimates of the time of transmission. We also identified situations when transmission time or direction was difficult to estimate by any method, particularly when transmission occurred long after the source was infected and when sampling occurred long after transmission. Applying our model to real HIV transmission pairs showed some agreement with facts known from the case investigations. We also found, however, that uncertainty on the inferred transmission time was driven more by uncertainty from time-calibration of the phylogeny than from the model inference itself. Encouragingly, comparable performance of the Markov time-slice model and the coalescent model—which make use of different information within a tree—suggests that a new method remains to be described that will make full use of the topology and node times for improved transmission time inference.

## Introduction

Identifying the time of infection is important for public health, clinical, and scientific reasons. For a population of hosts, i.e., on the epidemiological level, the timing of infections defines the incidence curve, which reveals temporal changes in infection in demographic or transmission groups, and can be compared against the timing of intervention strategies to assess their efficacy (Rosenberg and Biggar 1998; Williams et al. 2001; Quan et al. 2002; van Sighem et al. 2015). For one person who is newly diagnosed, identifying the time of infection would inform the relevant time window into the past for contact tracing. For a transmission pair, in which a known source infected a recipient, identifying the time of transmission can provide information on particular transmission circumstances and risks. For example, in mother-to-child transmission pairs, transmission can occur before, during, or after birth (Embree et al. 2000; European Collaborative Study 2005; Kourtis et al. 2006), and better knowledge of these relative probabilities could inform interventions (Newell 1998). In forensic cases, it is relevant to establish not only whether the accused infected the victim, but also whether this occurred at the time of the alleged crime or after the accused knew of their own infection (Abecasis et al. 2018; Ezeonwumelu et al. 2018). In an exclusive couple, knowing whether transmission occurred after few or many sexual encounters would inform the transmission probability per encounter (Wawer et al. 2005) and potentially allow comparison of transmission probabilities for different behaviors.

A variety of methods exist for inferring infection time of individuals. In some cases, the time of infection can be narrowed down to a period of likely exposure. During the acute infection stage, combinations of specific clinical markers can inform time of infection (Fiebig et al. 2003). Molecular biomarkers, such as CD4 cell count or virus sequence diversity, used singly or in combination, can provide an estimate of infection time (Giorgi et al. 2010; Skar et al. 2013; Giardina et al. 2019; Lundgren et al. 2022). When clonal virus sequences are available from the infected person, especially from multiple timepoints, the virus phylogeny can indicate the time of infection, at least under the assumption of a single founding genotype (Poon et al. 2011). When clonal virus sequences are available from the source as well as the recipient, and a time-calibrated phylogeny is constructed, the time of the most recent common ancestor of any source and recipient lineages is sometimes used as an estimate of the time of transmission (González-Candelas et al. 2013, for HCV), though more precisely this node provides an upper bound on how long ago the transmission event was (Leitner and Albert 1999; Leitner 2019). Such data have also been used with coalescent models that include the transmission event, in order to compare within- and between-host evolutionary rates (Vrancken et al. 2014), estimate the root age of a sample that spans hosts (Bletsa et al. 2019), or determine the transmission direction (Romero-Severson et al. 2017), but not yet for the goal of estimating the transmission time (though see also Lundgren et al. 2022, who used a coalescent model but obtained transmission time information primarily from biomarkers).

Here, we present a new model designed to ask how well transmission time can be inferred from phylogenies constructed from multiple HIV sequences representing the HIV populations in a source and recipient pair. We show that in the best-case scenario of fully-resolved and correct source-recipient trees, a simple model-based phylogenetic method can do quite well at recovering the time of the true transmission event. Working with real HIV data from known transmission pairs, we faced the challenge of accurately time-calibrating the trees, but we could still often recover transmission times that were consistent with known case histories.

## New Approaches

From a time-scaled virus phylogenetic tree (timetree) where two epidemiologically linked persons (hosts) were sampled, under the assumption that transmission was in only one direction, basic logical reasoning places bounds on the possible transmission time (Fig. 1A; Leitner 2019). The earliest transmission time cannot be prior to the most recent node that yields descendant tips in both hosts, because there would be no possible ancestral state assignments that could explain the data. It could be later, however, up until the first time by which both hosts have been sampled. We will call this logical, phylogeny-based time window for transmission the ‘phylogical window.’

**Figure 1:**
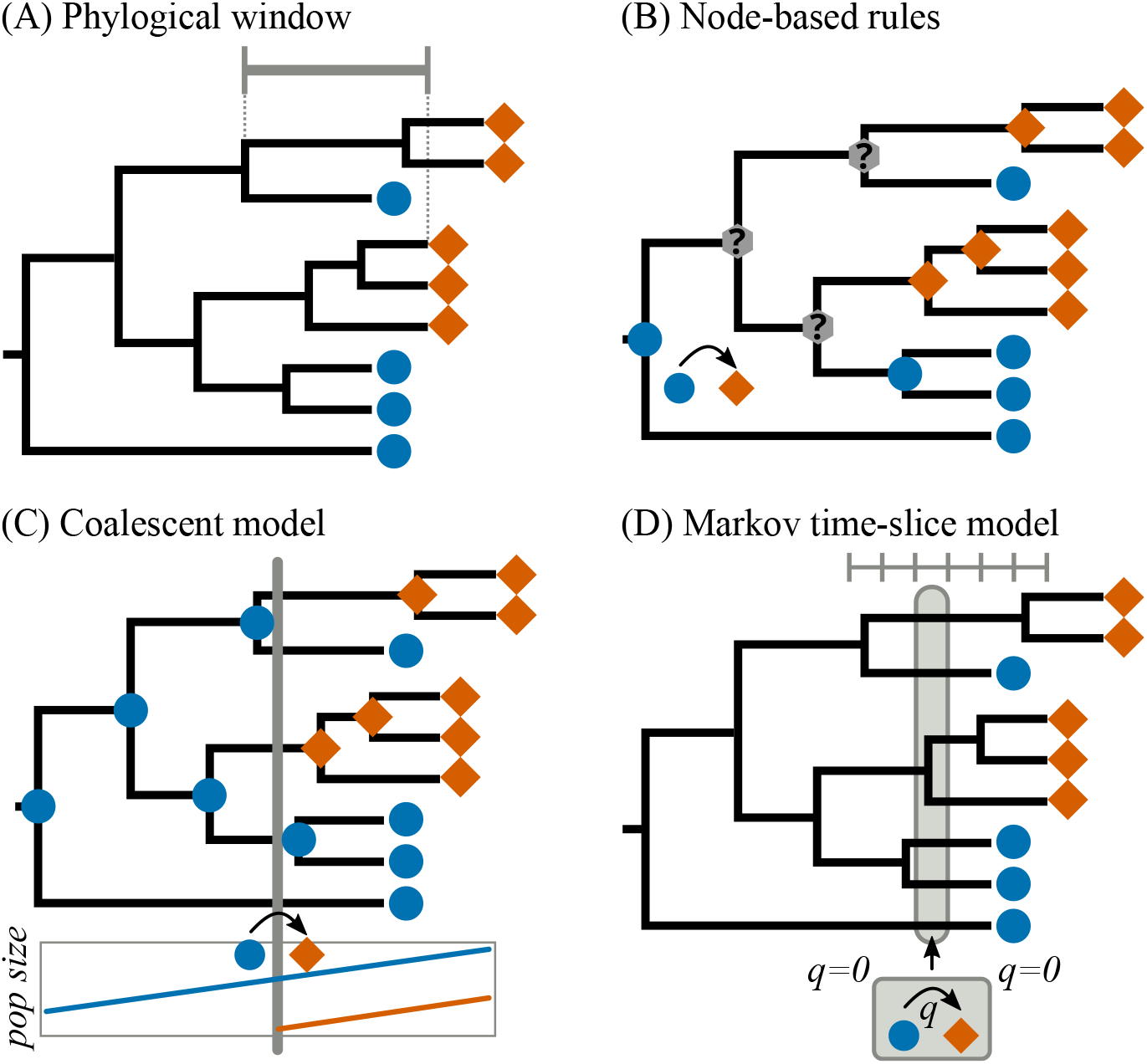
Methods for inferring transmission time and direction from a source-recipient phylogeny. In this small example, samples were obtained from one host at one time (blue circles) and from another host at two times (orange diamonds). The tree is time-calibrated. (A) The transmission time must logically be no earlier than the most recent node with descendants in both hosts, and no later than the earliest time by which both hosts have been sampled. This defines the phylogical window, shown by the gray bar. (B) Romero-Severson et al. (2016) defined rules for assigning each node as being present in one host (blue or orange) or equivocal (gray). The host state at the root node indicates the inferred source. (C) In a coalescent model, each node is first assigned to a host based on the transmission time, and the likelihood of each node depends on the virus effective population size at that time in the corresponding host. Here a linear function is shown for viral growth over time, with a bottleneck at the transmission event. (D) In our Markov model of transmission, the phylogical window is divided into time slices, one of which is assigned a rate of transmission (*q >* 0) from one host to the other. Statistical model comparisons are used to determine support for transmission occurring in each possible time slice and either possible direction.

Within this phylogical window, model-based approaches could potentially identify if any transmission times are more likely than others. In a coalescent framework, assuming a particular transmission time determines the host assignment for each node, and the overall tree likelihood is then computed from the dates of each node and used to estimate transmission time as a free parameter (Fig. 1C). Such models have been applied to source-recipient trees, but they inherently have many parameters (at a minimum, the infection times, transmission bottleneck sizes, and viral population growth rates in each host) and have proven unwieldy to fit (Romero-Severson et al. 2017). This type of model has also been used to aid timetree reconstruction when there are known boundaries on the possible transmission time (Vrancken et al. 2014).

Our goal was to determine whether a simpler model-based approach can improve transmission time inferences on HIV timetrees from persons who may have infected each other. Rather than treating the tree as a random draw from a model of the virus population history (including within-host dynamics as well as transmission, as the coalescent does), we simply view the host state as a label that evolves on a fixed tree. We use a Markov model with a single parameter *q* denoting the instantaneous rate at which lineages change from one host state to the other. This parameter serves merely to represent the stochastic nature of which lineage(s) happen to be involved in transmission, and we assume it is not affected by viral population dynamics. Under the assumption that transmission was only from one person to the other and at a specific time, we allow the transmission process to be active (*q >* 0) only in one direction and only during a particular slice of time (Fig. 1D). We use a slice of time with some width, rather than a single instant, to allow multiple lineages be transmitted. This is essentially a very specialized form of the phylogenetic model traditionally used for nucleotide evolution and other discrete-valued traits (Felsenstein 1981; Lewis 2001), additionally with the evolutionary rate parameters divided into epochs (Bielejec et al. 2014).

For statistical inference on a timetree, we fit many versions of the general model, each defined by the time slice and direction in which transmission is possible. Slices can be defined either as arbitrary intervals (e.g., one month wide, as we did for our simulation studies) or based on known events (e.g., a known period of contact between the source and recipient). The best-supported model then provides a single answer about who infected whom and when. Additionally, we can incorporate uncertainty in this inference. We compute the support for each model version—each transmission time and each transmission direction—and then summarize across models, weighting each by the evidence for it.

Hence, our model yields not only an estimate of the transmission time, but also of the transmission direction. Compared with previous work that used a Markov model for this purpose, we represent transmission more realistically, as a brief event rather than a continuously-ongoing bi-directional process (Villabona-Arenas et al. 2022). Another method is to work from the tips back to the root and assign each node an inferred ancestral state based not on a model, but on a set of parsimony-inspired rules (Fig. 1B; Romero-Severson et al. 2016, 2017). This has the advantage of being very simple and requiring only a tree topology, not branch lengths or time calibration. Because it is heuristic rather than statistical, however, there is no basis for reporting confidence in the inference, and features such as more tips or earlier sampling time for the recipient than the source can mislead the inference (Romero-Severson et al. 2017). We compare our time-slice model inference of transmission direction with this node-based rules method.

## Results

### Simulation tests

We first used simulations to assess how well our time-slice model can identify the true time and direction of transmission. We considered 23 different combinations of infection and sampling times (Fig. 2A). These histories encompass long and short times between infection events and between infection and sampling, and sampling of the source and recipient once each in either order. In the simulations, the source infects the recipient at a single point in time, when multiple virus lineages could be transmitted. For each history, we generated 50 replicate trees by coalescent simulation, each with 20 tips per host; one representative tree for each history is shown in Fig. S1. This coalescent model assumed that the virus effective population size within a host increased linearly with time, as has been empirically documented for HIV (Shankarappa et al. 1999; Zanini et al. 2015). Note that the generating model was thus more complex and realistic than the inference model, which ignores virus population dynamics.

**Figure 2:**
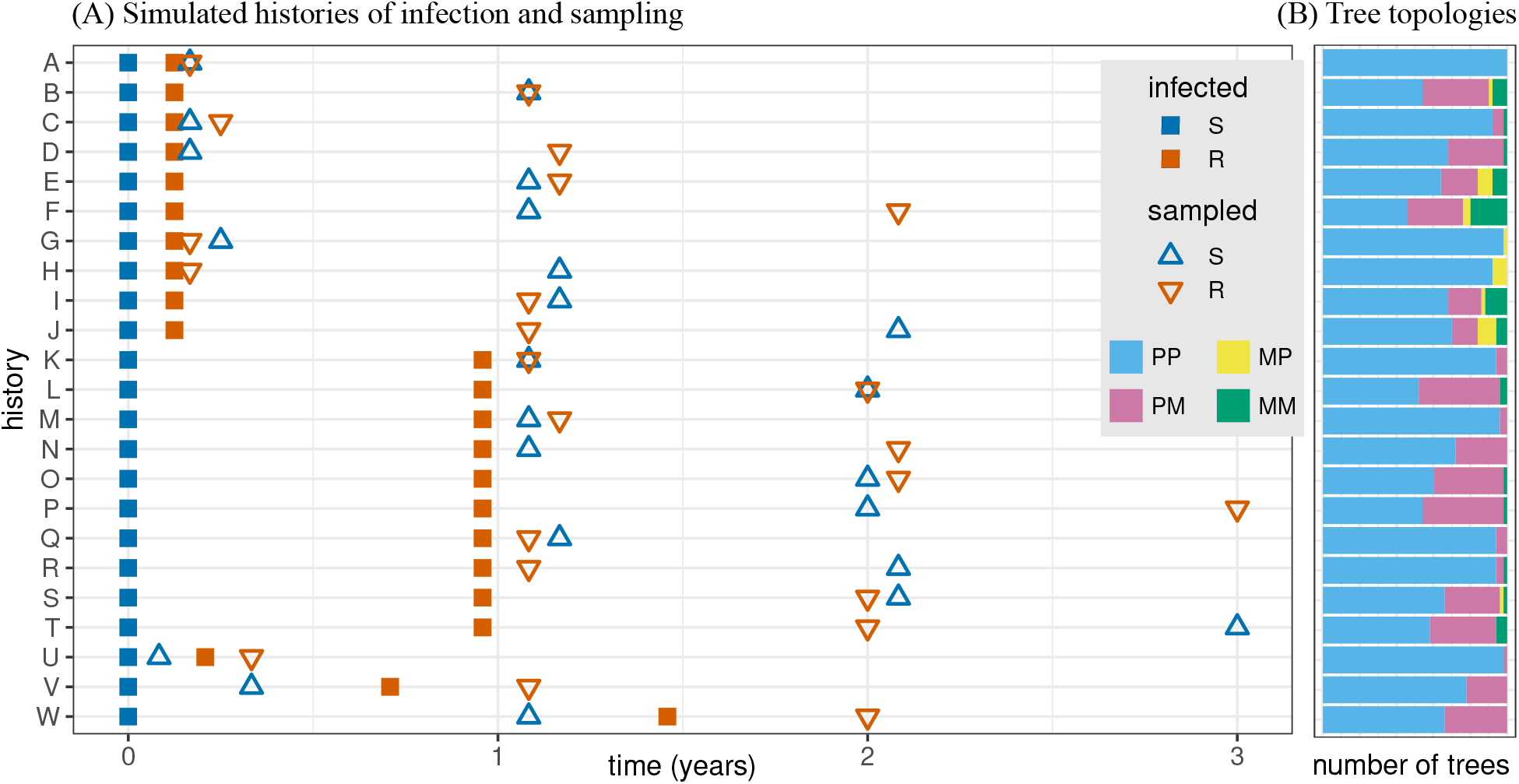
Data were simulated under 23 different combinations of infection and sampling times for the source (S) and recipient (R). (A) Transmission from source to recipient could occur soon (1.5 months, histories A–J) or later (11.5 months, K–T) after the source is infected. The source could be sampled at the same time (A–B, K–L), before (C–F, M–P), or after (G–J, Q–T) the recipient. Additionally, if treatment is not effective, it could be possible that transmission to the recipient occurs after the source is diagnosed and sampled (U–W). (B) For each history, most trees are para/polyphyletic for the source and recipient (PP), though some instead are monophyletic for the recipient (PM) or for the source (MP). Trees with monophyletic clades for both the source and recipient (MM) sometimes occur when the sampling times are much later than the infection times.

We aimed to analyze trees on which the virus sequences from the two hosts are interdigitated. This tree topology is not universal in direct transmission pairs, but it is common [Leitner and Romero-Severson (2018) report it for 52% of direct pairs]. Furthermore, on such topologies the transmission direction is often not revealed by the node-based rules method of Romero-Severson et al. (2016, 2017). In the simulations we therefore founded the infection in each host with a relatively high level of initial virus diversity, producing trees that were mostly para/polyphyletic for the source and recipient (Fig. 2B). The number of distinct recipient clades on a tree defines the minimum number of transmitted lineages required to explain the sampled data; this number was never more than 8 in our simulations, and typically 2–4 (Fig. S2A), in agreement with empirical trees for direct transmission (Leitner and Romero-Severson 2018). The transmission time inferred by the time-slice model did not depend on the minimum number of transmitted lineages for most histories, though for some the accuracy improved slightly as more recipient clades were sampled on the tree (Fig. S2B).

#### Model inferences of transmission time improve on the phylogical window

Because our goal with simulation testing is to assess the overall power and accuracy of our time-slice model, we summarize results across all trees within each history rather than consider each tree separately. For every simulated history, the 90% credibility interval (CI) of the time-slice model-based transmission time inference, summed across trees, included the true transmission time (Fig. 3). Additionally, the single best-supported month-long time slice contained the true transmission time for 17 of the 23 histories. The 90% CI was also narrower than the phylogical window for all histories that spanned more than four months. When one or both hosts were sampled long after the transmission event, the phylogical window could not rule out a recent transmission event but, encouragingly, our model could: on all of the 16 such histories, the model 90% CI ended at least five months before the final sampling time (Fig. 3, transmission histories B, D, E, F, H, I, J, L, N, O, P, R, S, T, V, W).

**Figure 3:**
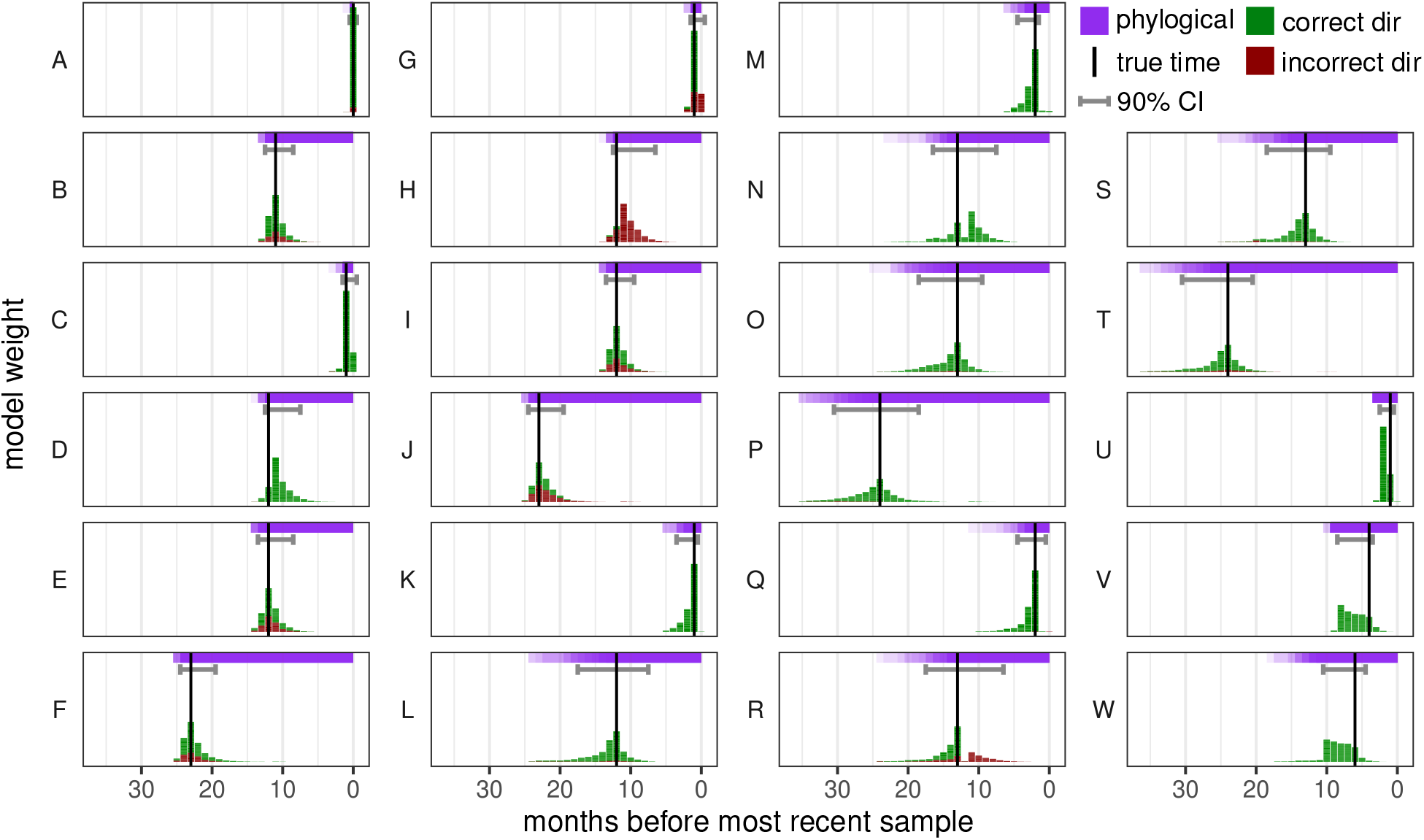
Inference of transmission time and direction on simulated data using our Markov time-slice model (Fig. 1D). Each panel shows results for one of the 23 transmission histories (Fig. 2A), with the true time of transmission marked by a vertical black line. Histogram bar heights show the inferred model weight for the transmission event occurring at each time slice, summed over 50 tree replicates. Results for one tree may span multiple time slices if there is uncertainty in the inferred time. Gray horizontal intervals show the 90% credibility interval for the time-slice model-inferred transmission time, for the model weights summed across trees. The histogram bar fill color shows whether the model had transmission in the correct direction (green) or the incorrect direction (red). The phylogical window is also shown for each history (horizontal purple band), with variation among the replicate trees indicated by transparency.

The beginning of the phylogical window is a good estimate of the transmission time when the recipient is infected soon after the source (histories A–J, U), even if sampling was much later, because the most recent node with descendants in both hosts tends to fall deep in the tree. Otherwise, the phylogical window extends much deeper in the tree than the true transmission time (histories K–T, V–W). Comparing the model’s single best-supported slice with the start of the phylogical window, they were the same on 39% of the 1150 total trees, and our model estimate was closer to the true transmission time on 52%. The time-slice model performed worse in only three histories, those in which one host was sampled soon after transmission and the other was sampled long after (histories D, H, N; discussed further below).

#### Uncertainty in transmission time caused by time between infection events and delays until sampling

Uncertainty in the inferred transmission time is greatest when two conditions are both met: a long time from infection of the source until infection of the recipient, and a long time from the transmission event until sampling one or both hosts. Histories P and T provide the most extreme version of these events and have among the largest transmission time credibility intervals (Fig. 3).

The influence of the time between infections is apparent from comparing the smaller CIs of histories F and J (short time from source infection to recipient infection) against the larger CIs of histories N and R (same sampling times but longer time until recipient infection). On trees in the latter histories, clades from the second-sampled host fall on long branches extending beyond the first sampling time, providing little information to distinguish when transmission occurred along these branches (Fig. S1).

Interestingly, considering the effect of the duration of time from transmission to sampling, sampling both hosts at the latest time produces no more uncertainty—or even a bit less—than sampling one host at an earlier time. Compare, for example, the CIs of histories E, I, O, and S against the equivalent or larger CIs of the histories that differ only in sampling one host earlier (respectively, D, H, N, and R). On trees in the former histories, the inferred transmission time is both more precise and more accurate because more of the time slices contain branches from within both hosts, better informing the model’s transmission parameter. Alternatively, when both hosts are sampled soon after the transmission event, the transmission time inference is much more precise (smaller CIs in histories K, M, and Q).

#### Delay between recipient and source sampling misleads transmission direction more than transmission time

In addition to inferring transmission time, our time-slice model also infers the transmission direction. Across our simulated histories, the model performs similarly to the node-based rules (Fig. S3).

We expected that sampling the recipient before the source would make it difficult to infer the correct transmission direction, because older lineages would be more likely to be captured in the recipient’s sample. This turned out to be only partly true. Histories H and J, which have a long time between recipient and source sampling, were the most challenging for both methods. In these histories, the recipient’s sampled sequences closely resemble the source’s sequences at the time of transmission, and the later-sampled source sequences tend to appear as descendants from them (Fig. S1). Histories R and T have the same sampling structure but a longer time period between source and recipient infection. These histories presented milder problems for our time-slice model inference, and the node-based rules method performed well on both. When the source is sampled soon after the recipient, the inference of transmission direction by either method is no worse than when the recipient is sampled soon after the source (compare E vs. I, and M vs. Q).

Even when the model is more than 25% wrong about transmission direction (histories E, F, H, I, J, R), its transmission time estimates are generally not far off, as discussed above (Fig. 3). Most extremely, for history H the model places the transmission time shortly after the recipient is sampled, causing it to report confidently but incorrectly that the recipient infected the source. The true transmission event was only slightly before the recipient was sampled, however, so the error in the transmission time is not large. (This qualitative error, however, of inferring a transmission time that is after rather than before a sampling time is seen also in histories D and N.) In history R, the model provides substantial weight for two alternative scenarios: transmission in the correct direction at the correct time (or slightly earlier), versus transmission in the wrong direction at a later time (after the recipient is sampled). Therefore, if either the transmission time or the transmission direction were known from external information, the other would be inferred correctly by the model.

### Empirical case studies

As our simulation tests established that our time-slice model could infer the time of transmission for ideal data, we also wanted to test how well it would perform on real-world data. Not only does real data evolve according to very complex evolutionary conditions, typically much less certainty exists about transmission direction and, in particular, the exact time of transmission. Thus, from a large compilation of known HIV transmission pairs (Leitner and Romero-Severson 2018), we selected three that had the best potential for estimating informative time-calibrated phylogenies (many clones per host, multiple sampling times), presented a challenge for inferring transmission direction (para/polyphyletic topology reported in previous work), and spanned a variety of transmission modes.

The first case study was a mother-to-child transmission pair (Simonich et al. 2016, hereafter ‘MTCT pair’), with three sampling dates for the mother and two for the child. The child tested HIV negative 8 days after birth and then HIV positive 114 days after birth. The second case study involved sexual transmission between two men (Frange et al. 2013, hereafter ‘MSM pair’). The recipient was infected approximately 37 days before being sampled, and the source was sampled 2 weeks later. The third case study was HIV transmission during a robbery at knife-point (Kao et al. 2011, hereafter ‘Knife pair’), which is an unusual transmission mode but has the advantage that the exact day of transmission was known. The sequence data included a sample from the source 225 days after transmission and a sample from the recipient 19 days after that.

For each case study, we inferred a time-calibrated phylogeny from the archived sequence data, used our transmission time-slice model to estimate the time of transmission from that tree, and validated the findings against the known facts of the case history. Ideally, the dated phylogeny would be constructed based only on the HIV sequences from the two hosts and tip sampling times. We refer to these analyses as transmission ‘direction-naive’ analyses. In some cases, however, we found that such a tree contradicted facts of the case, yielding a phylogical window that excluded the known transmission time. We suspected that this contradiction arose from a combination of local and global evolutionary rate variation across the virus tree (Wertheim et al. 2012; Worobey et al. 2014; Bletsa et al. 2019), which is not accounted for in standard molecular clock models. We therefore additionally made trees that included information about the transmission direction in the tree inference model by allowing molecular clocks to run separately in the two hosts, using the method of Vrancken et al. (2014). We refer to those analyses as transmission ‘direction-aware’ analyses. A byproduct of making such direction-aware phylogenies was an estimate of the transmission time from a coalescent model, which we could then compare against our time-slice model estimate.

#### Direction-naive HIV phylogeny provides plausible time of transmission (MTCT pair)

For the mother-to-child transmission pair the time of infection was not known, but from the case history we could define a clinical window for transmission extending from 3.8 months after birth (the date of the first HIV positive test of the recipient), back to 5–6 weeks before birth (allowing that an HIV test may still register negative soon after infection; Delaney et al. 2016). When we constructed a phylogeny without information on transmission direction and applied our time-slice model, the most likely time of infection was during the month of birth, and the 90% CI of transmission time extended from then back two months (Fig. 4). Our time-slice transmission time inference is thus partially consistent with the clinical data, though it is inconsistent with the proposition in the original paper that (based on the test dates) transmission was during breast feeding. The phylogical window did not argue against transmission after birth because it inherently extends until the date of the recipient’s first HIV positive test. It did, however, allow that transmission could have been even earlier, with the 90% CI of the beginning of the phylogical window extending 2.7 months before birth to 1.3 months after. Our time-slice model thus provided a tighter estimate of transmission time than the phylogical window did.

**Figure 4:**
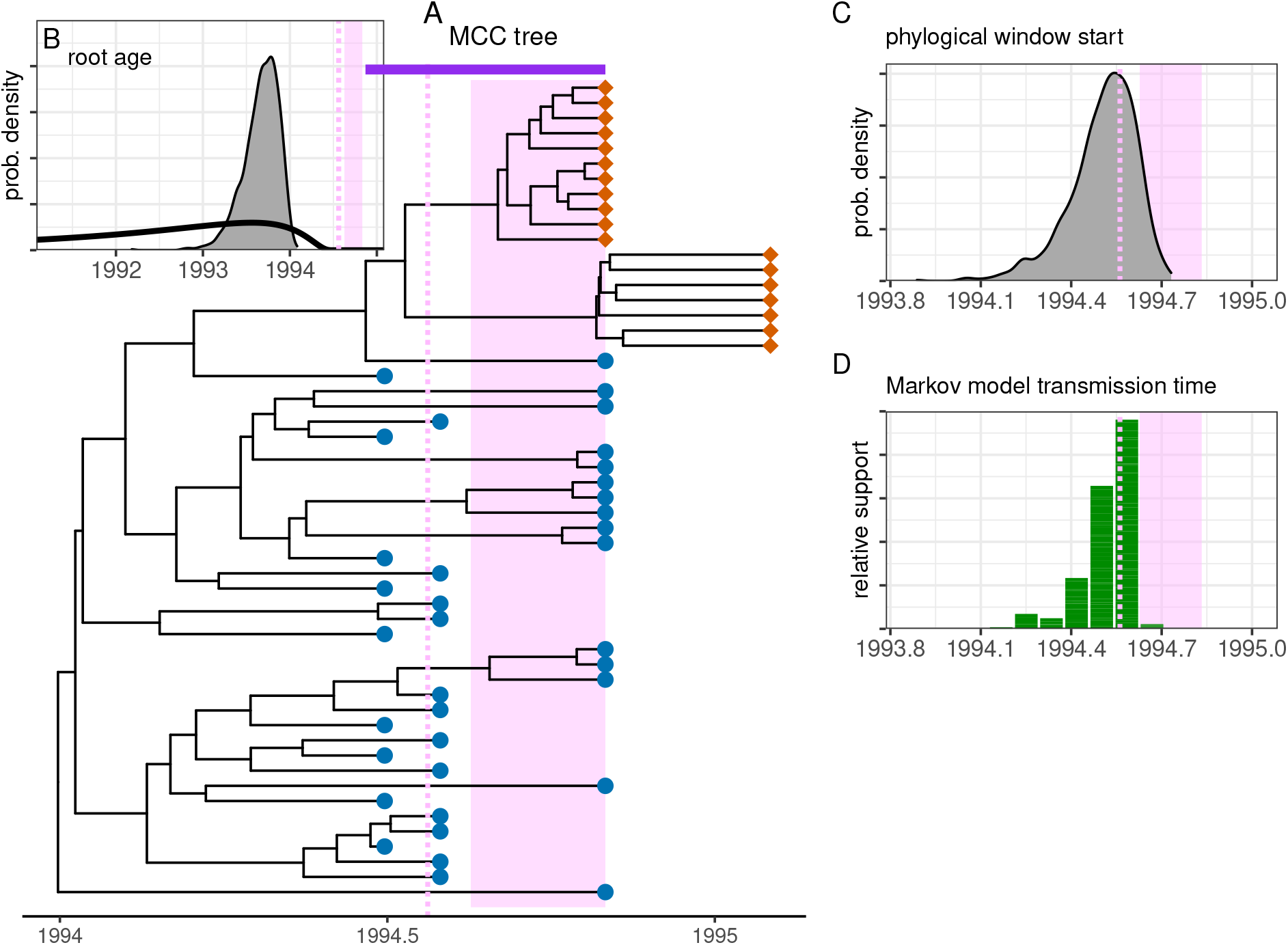
MTCT pair case study. In all panels, the pink vertical line marks the date of birth, and the shaded pink rectangle marks the window between the recipient’s last HIV negative test and first HIV positive test (though this window is not a definitive range of possible infection times because an HIV test may still register as negative soon after infection). (A) Maximum clade credibility tree, inferred with direction-naive methods, with its phylogical window marked by the horizontal purple bar. (B) Root age for Bayesian inference of the tree. (C) The phylogical window’s start across the posterior set of trees. In (B) and (C), posterior distributions are shown with shaded gray areas, and the prior distribution in (B) is shown with a thick black line. (D) The time-slice model-inferred transmission time (one histogram bar per slice) and direction (all green, meaning correct direction).

Using instead the direction-aware phylogeny suggested that the inferred transmission time was even earlier (Fig. S5; 90% CI of 2–7 months before birth using our time-slice model, 0.2–5.1 using the coalescent model, and 1.7–7.1 using the start of the phylogical window). These estimates seemed rather early, considering the HIV negative test 8 days after birth. The inferred transmission direction was correct on all posterior trees, for both the node-rules and time-slice methods.

#### Direction-aware phylogenetic reconstruction improves transmission time inference (MSM pair)

For the MSM pair, the direction-naive phylogeny led to an inferred transmission time that was significantly later than the date estimated by the original study based on symptom onset and laboratory tests (Fig. S6): the 90% CI of the transmission time was 10–50 days later than the reported date, and the phylogical window excluded the reported date with 100% support. Additionally, the time-slice model incorrectly placed the transmission event after the recipient’s sampling time, and consequently inferred the wrong transmission direction, with 66% support.

Using instead the direction-aware phylogenetic reconstruction, we could infer a substantially earlier transmission time. Hence, defining whether each tip came from the source or recipient, but with an uninformative prior on the transmission time, the model-estimated transmission time had a 90% CI of 10–110 days before the clinically-determined infection date of the recipient using our time-slice model, and 0–115 days before using the coalescent model (Fig. 5). The start of the phylogical window had a 90% CI of 11–157 days before the clinically-determined infection date. Interestingly, placing a strong prior on the transmission time had little additional effect (90% CI 10–100 days before from our time-slice model, 4 days after – 100 days before from the coalescent model, and 8–162 days before for the start of the phylogical window) (Fig. S7). Both our time-slice model and the node-based rules method obtained the correct transmission direction with 100% support; note that the correct transmission direction was provided as an input to the coalescent model.

**Figure 5:**
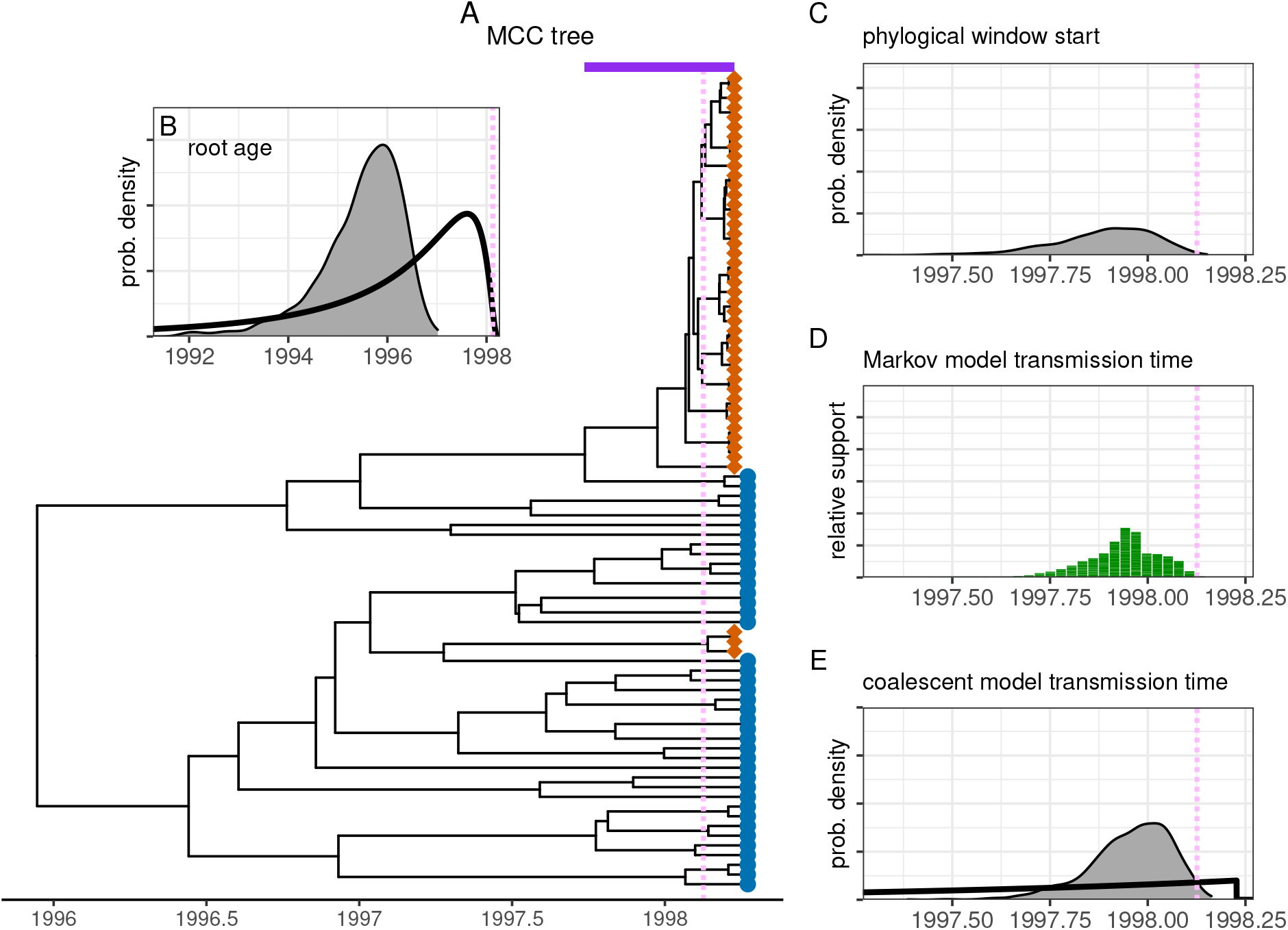
MSM pair case study. These results are from direction-aware phylogenetic inference, with an uninformative prior on the transmission time (thick black line in panel E). The pink vertical line in all panels marks the transmission date estimated by the original study. Figure components are otherwise as in Fig. 4, with (E) additionally showing the posterior distribution of transmission time derived from the coalescent model used for tree inference.

#### Uncertainty in time-scaling trees inhibits precise transmission time inference (Knife pair)

For the Knife pair, the transmission date was known precisely from case information, but the direction-naive reconstruction suggested an inferred transmission time that was too late because the phylogical window excluded the true transmission time with 100% support (Fig. S8). The transmission direction from our time-slice model was correct, however, with 100% support.

The direction-aware reconstruction, instead, provided transmission times consistent with the true date (Fig. 6). When no additional information on the transmission time was provided, the phylogical window included the true transmission time with 82% support, and the true transmission time fell near the center of the time-slice model inference (39% quantile) as well as the coalescent model (48% quantile). The transmission direction was correctly inferred by the node-based rules (98% correct, 2% uncertain) and our time-slice model (100% support).

**Figure 6:**
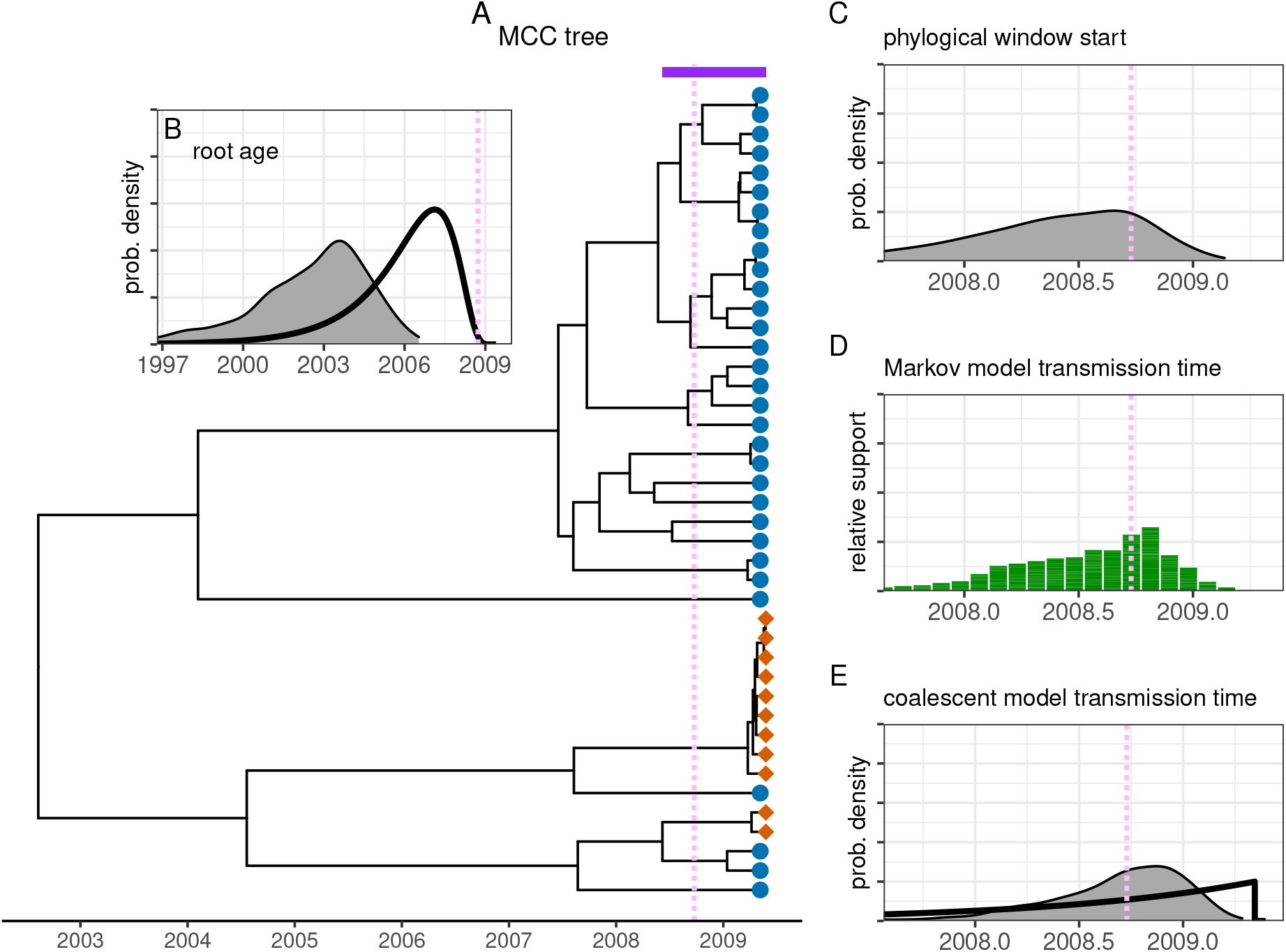
Knife pair case study. These results are from direction-aware phylogenetic inference, with an uninformative prior on the transmission time. The pink vertical line marks the known date of transmission, obtained from the case information. Figure components are otherwise as in Figs. 4 and 5.

Our time-slice estimate of time of transmission had a broad CI that could be due to (at least) two factors. First, the uncertainty due to time calibration problems can be quantified by variation in the single best-supported time slice among the posterior trees (e.g., its 90% CI). Second, the uncertainty due to model inference can be quantified by the typical size of a credibility interval on one tree (e.g., the median of the 90% CI width from each tree). We found that the former quantity is larger than the latter—14 vs. 6 months—meaning that the overall uncertainty in transmission time was set more by time calibration than by our model inference on the tree.

If instead the phylogeny was additionally inferred with knowledge of the true transmission time (strong prior with 90% CI width 3.9 months), transmission time inference closely followed that prior for the coalescent model (90% CI width 3.6 months), and was broader for our time-slice model (90% CI width 10 months) (Fig. S9).

## Discussion

Our goal was to assess how well the time of a transmission event can be obtained from a phylogeny based on multiple virus sequences from both the source and recipient hosts. We developed a Markov transmission time-slice model, which is simpler (with only one free parameter) and easier to fit than existing coalescent models. Simulating many transmission and sampling histories showed that our time-slice model generally correctly inferred the known time of transmission, though with uncertainty. When testing our time-slice model on empirical data, however, we found that including information about the transmission direction was sometimes necessary to obtain trees that were logically consistent with the known case history. In this process, we found that the transmission time obtained by our simple time-slice model was comparable to that obtained by the computationally more costly coalescent model used in tree construction. Because these two modelling frameworks make use of somewhat different aspects of the phylogeny, we suggest that future development of a model combining their properties could provide better inference of transmission time.

Our simulation study showed that phylogenies with interdigitated tips from the source and recipient (known as para/polyphyletic topology; Romero-Severson et al. 2016) contained substantial information about transmission time, which could be extracted with our time-slice model. Without a model, the phylogical window can not rule out transmission times that are too late, and it also allows transmission too early particularly when the recipient was infected long after the source. Our time-slice model-based results were substantially better in both of these aspects. The most challenging scenarios for transmission time inference were those in which much time had elapsed from the source’s infection to the recipient’s infection, and then to the sampling of either host. This finding is in line with previous results that showed that inferring transmission order is more difficult when longer times are involved (Romero-Severson et al. 2014). We also found that transmission time could be inferred accurately even when transmission direction was not, such as when the recipient was sampled long before the source. Additionally, in some scenarios, knowledge of the true transmission direction would enable a more accurate estimate of the transmission time. When the goal is only to infer transmission direction, not time, however, the simpler node-based rules method (Romero-Severson et al. 2016, 2017) generally performed as well as our time-slice model.

In our empirical applications, for one transmission pair (MTCT pair) were we able to infer a informative phylogeny from the sequence data alone, and then infer a reasonable transmission time from that. For the other two pairs investigated, transmission times—whether from the time-slice model or the phylogical window—were consistent with case information only when the trees were built with a prior on node times that was informed by who was the source and who the recipient (direction-aware reconstruction). The method used for constructing such trees yielded a coalescent model estimate of the transmission time, while the time-slice model takes only the posterior trees as input and yields a separate estimate of the transmission time. While this is not logically circular (because the time-slice model does not make the coalescent assumptions that go into constructing the tree), it does show that inferring transmission time from trees made with generic, direction-naive methods may often be misleading. Indeed, previous work has shown the importance of accounting for different evolutionary rate distributions on different portions of a virus phylogeny (Worobey et al. 2014; Vrancken et al. 2017; Bletsa et al. 2019). Thus, our exploration raises two promising avenues for future work.

First, the similarity in transmission time estimates from the Markov time-slice and coalescent models—both in value and uncertainty—on the same trees suggests that the former may be sufficient for co-inference of tree and transmission. One advantage of using the time-slice model instead of a coalescent model for this purpose is its simplicity: our version has only one free parameter and makes no explicit assumptions about the dynamics of viral effective population size within each host. More importantly, the time-slice model may extend more flexibly to clusters in which the pairing and timing of transmission events is unknown. Existing coalescent models can take pre-specified sources and recipients (Vrancken et al. 2014) or can piece together infection chains from windows of plausible transmission times (De Maio et al. 2016; Lundgren et al. 2022). However, they do not yet consider the dynamics of cluster formation as part of the model, allowing, for example, that transmission is more likely between people with the same exposure modes. Rather than the statistical model comparisons used in the present paper, an expanded Markov time-slice model could be cast with a free continuous-valued parameter for the start time of each active slice. With the host label evolving stochastically on the tree, this becomes analogous to the approach of Landis (2016), in which a phylogeny is co-inferred with the biogeographic history of the clade. Divergence time estimation is there informed by a paleobiogeographic model, which allows dispersal as a stochastic process within a set of specified geologic constraints. The analog here is a model of a contact network, within which transmission occurs as a stochastic process under a set of constraints imposed by any case history knowledge. At best, this could provide a means for co-inferring phylogeny, transmission times, and transmission pairs within a cluster. Even if such estimates were not precise, however, the approach could yield statistically robust inference of the incidence at the epidemiological level.

Second, we propose that neither the coalescent nor the Markov time-slice model is ideal for transmission time inference (or co-inference) with a phylogeny, and that therefore a more powerful model is waiting to be formulated. The coalescent model describes the viral growth dynamics within each host, using that to draw information from the times of all nodes on the tree. It does not, however, describe the dynamics of the transmission event itself. The inferred number of lineages transmitted and their relationships thus do not affect the coalescent likelihood directly (only indirectly, through the consequent host identities of each node). The Markov time-slice model take the opposite perspective. It does not model viral growth dynamics, and hence ignores information from node times outside the active slice, but it does model the transmission event as a dynamic process. Because the within-host dynamics and transmission dynamics each evidently provide information about the transmission time—evidenced by efficacy of both the coalescent and time-slice models—we propose that modeling both together would provide more power. We believe that such a model cannot be trivially constructed by multiplying the coalescent and Markov time-slice likelihoods together, because the two likelihoods do contain some of the same tree information. Including a coalescent probability for each node in the pruning algorithm, or including a small window of unidirectional migration in the coalescent model, both seem like plausible approaches, but they make different assumptions about the lineages that are transmitted. We therefore speculate that the unified model may best be formulated by building from slightly more detail about the biology of the transmission process. It remains to be seen if an improved model can be defined, and if it does indeed provide more power to infer transmission time from viral phylogenies. Recent work defining how likelihood can be computed for more explicit forward-in-time transmission models (King et al. 2022) could provide the theoretical framework to bridge the modeling approaches we have discussed in this paper.

## Materials and Methods

We apply both existing and new methods for inferring the transmission time and direction from a phylogenetic tree. Overall, we assume that a phylogeny has already been obtained, with multiple clonal virus sequences from the two people in a transmission pair. We further assume that transmission was in only one direction, from the source to the recipient; we do not assume that the transmission direction is known, but we show how this information could be included in our model if it is available.

### Heuristic approaches

Using the tree topology only, without branch lengths, Romero-Severson et al. (2016) introduced a simple set of rules to infer which host was the source. Working from the tips back toward the root, each node is assigned an identity according to the identity of its two daughter nodes or tips (Fig. 1B). If both daughters are in the same state, that is the assigned node state. If the two daughters hold different host states, the assigned node state is uncertain. If one daughter is uncertain and the other holds a host state, the node is assigned that host state. The source is the host inferred at the root node. This node-based rules procedure often, but not always, produces state reconstructions that are the same as parsimony under the requirement that state changes are only in one direction. It is very quick to apply and does not require a time-calibrated tree. It does not have a clear statistical basis, however, so it is not possible to infer confidence in the answer.

Using a time-calibrated tree, Leitner (2019) explained how the node dates provide constraints on when transmission could possibly have occurred (Fig. 1A). Specifically, the transmission time cannot be prior to the most recent node that yields descendant tips in both hosts, and it of course must be before the first time by which both hosts have been sampled. Simple reasoning provides no further information on when in this ‘phylogical’ time window transmission occurred. Here, our model-based approach aims to be more precise.

### Markov time-slice model

For inferring transmission time and direction, we use a simple phylogenetic Markov model that ignores virus population dynamics in each host. The host label is treated as evolving on the fixed tree in the same manner as a nucleotide state or other discrete-valued trait (Felsenstein 1981; Lewis 2001). The single parameter *q* is a rate at which lineages change from one host state to the other. We allow this transmission process to be active (*q >* 0) only in one direction and only during a particular interval of time, a ‘time slice’ (Fig. 1D). For our simulation study, the intervals were pre-determined by slicing the duration of the tree into equal widths. Slicing started at the latest sample (so the first slice ends at the most recent tip) and worked back toward the beginning of the phylogical window (whose slice may therefore be somewhat smaller). For empirical analyses, the slices can be adjusted to test specific hypotheses, such as whether transmission was before or after some event from the case history. If it were suspected that the transmission process were ongoing via numerous contacts, the model could be arranged so one slice spans more or all of tree (and modified to allow bidirectional transmission if necessary). Our model requires a time interval for transmission, rather than a single moment, because the parameter is an instantaneous rate; the probability that a given lineage in the host is transmitted to the source is *qdt* for an infinitesimal time interval *dt*. Thus, using a slice of time allows multiple lineages to be transmitted (otherwise, a Markov model would by definition not allow multiple lineages to change host state at a single moment of transmission). We expect smaller values of *q* when fewer lineages are transmitted, but the parameter has no specific physical interpretation, and we also expect smaller values of *q* when the time slice is wider if transmission was during a single contact.

We fit many versions of our model to each dataset, with each version defined by the direction and time slice of transmission. For example, in Fig. 1D, the model version depicted has transmission during the fourth time slice (counting back from the most recent tips) and in the direction from the blue host to the orange. A total of twelve model versions would be fit to that tree, to include each of the two transmission directions for each of the six time slices. For fitting each model version, the likelihood of the host tip states given the model and the tree is obtained via the pruning algorithm (Felsenstein 1981, with epochs as in Bielejec et al. 2014). Briefly, referring to the Fig. 1D example, we begin with the known host state of each tip. We first walk back on the tree from the tips to the more recent edge of the active time slice. The host state must remain unchanged along these branch portions. Then across the active time slice, for each lineage ending in the recipient host state, we compute the probability that the lineage began in the source host state at the older edge of the time slice; this depends on the rate parameter *q* and the width of the time slice. Lineages ending in the source host state began in the same state with probability one, due to the model’s assumption of unidirectional transmission. Finally, these probabilities are propogated back to the root with no further state changes allowed if there are no additional active time slices. At each node along the tree, probabilities are combined so that the parent state is the same as the two daughter states. When the model version specifies the transmission direction is from, say, the blue host to the orange (Fig. 1D), the root state must logically be blue when combining the conditional probabilities at the root (Goldberg and Igić 2008).

We used Bayesian inference for our model-fitting. The prior on the single parameter, *q*, was assigned to be exponential with rate 1/year for a time-slice width of one month. This is a somewhat arbitrary choice, but it reflects the empirically-supported expectation that the transmission bottleneck is more likely to be smaller than larger (Joseph et al. 2015; Kariuki et al. 2017). A prior supporting larger values of *q* could push the inferred transmission time closer to the tips of the tree by allowing more lineages to change host state within the active time slice. We found, however, that doubling or halving the prior rate for *q* had little effect on the results across the simulation histories (Fig. S10). Because *q* is a nuisance parameter (in that its value does not provide answers to our questions about transmission time or direction), we do not report fitted values for it. Indeed, we do not estimate it at all: instead, we computed the marginal likelihood of each model version by directly numerically integrating the posterior distribution over all possible values of *q,* [0, ∞). We then compared the statistical support for each model version, defined by its posterior probability, which is the product of its marginal likelihood and its prior probability. Our simulation study assumed equal prior probabilities for each model version, but if a case history provided informed priors on transmission time or direction, that information could be incorporated at this stage. To compare model support for the two transmission directions, we added together the weights for all models with one direction (across all time slices), and all models with the other direction (e.g., Fig. S3B). Alternatively, we can express our confidence in the inferred direction by computing the Bayes factor for support of one direction over the other as the ratio of their marginal likelihoods (again summed across time slices). We consider a Bayes factor of 3 as positive support for the favored model; if less, the inferred transmission direction is uncertain (Kass and Raftery 1995) (e.g., Fig. S3C). Similarly, we computed the support for transmission in each time slice as the total weights of the models that have that slice active (summed over transmission direction). Code for fitting our model is provided at https://github.com/MolEvolEpid/timeslice.

Other model structures are possible to capture similar biological detail. For example, instead of pre-defining non-overlapping time slices, one could instead give the model a free continuous-valued parameter for the active slice start time. This would change the statistical inference problem from model comparisons to parameter estimation, at least for transmission time. The latter is often an easier proposition, but in our case the former is also easy because we can efficiently compute the marginal likelihood. Alternatively, one could formulate a model in which multiple lineages may change host state at the same moment, with continuous-valued parameters for the time of the event and the proportion of lineages transmitted. We prefer our time-slice formulation, however, because it has only one free parameter, and it provides flexibility for situations in which transmission was concentrated in time but not instantaneous, such as from mother to child, or through repeated contacts with the same sexual partner.

### Simulation tests

To assess how well our Markov model could identify the true time and direction of transmission, we simulated idealized data under a variety of transmission histories (Fig. 2) designed to test whether inference is more challenging in circumstances that may erode the phylogenetic signal of recipient tips nested within source clades. We therefore particularly included situations when infection of the recipient was soon after infection of the source, when sampling was long after infection, and/or when the recipient was sampled before the source.

Under each transmission history, we generated 50 trees by simulating a continuous-time coalescent process in which the rate of coalescence in a host depended on the viral effective population size in that host. Beginning from the infection time, the simulator assumed linear growth in viral effective population size in each host, following data for HIV (Shankarappa et al. 1999; Zanini et al. 2015). Coalescent probabilities under this growth model were derived by Romero-Severson et al. (2014, 2017). For this linear growth function, we assumed that the infection of each host was founded with an initial effective population size of *α* = 20 (where *α* = 0 means transmission of a single lineage, as the rate of coalescence in the recipient becomes infinite at the moment of transmission, and *α >* 0 means that transmission of multiple sampled lineages is possible).

This function also assumes that the effective viral population size within each host grew at a rate of *β* = 3 per generation, using a within-host generation time of 1.5 days. Transmission to the recipient was assumed to occur at a single moment. For each simulated tree, we sampled 20 tips at one time for the source and for the recipient. These formed the trees on which we performed our inference; examples are shown in Fig. S1. Note that on each tree, the number of distinct recipient clades is substantially less than *α* = 20 (Fig. S2A). The number of distinct recipient clades is the minimum number of transmitted lineages needed to explain the tree, but the actual number of transmitted lineages may be larger if multiple lineages within one clade were transmitted. The coalescent simulation code is available at https://github.com/MolEvolEpid/biophybreak.

In these simulated data, the trees are by definition correct and fully resolved, without uncertainties or errors arising from stochastic sequence evolution or phylogenetic inference algorithms. They represent a best-case scenario for the data in that sense, although they still have only an empirically-reasonable number of tips. Note also that our simulated trees include virus population dynamics, while our inference model does not. We are thus testing, to some extent, whether the simple Markov model performs reliably on data generated under a more complex, realistic process.

Additionally, we performed a rough test of whether our simulated trees could be recoverable under a realistic mutation rate for HIV. We selected two histories, C and N, to represent shorter and longer timespans in our simulations. On each tree in those histories, we simulated sequences, inferred a timetree from those sequences, and then fit our time-slice model. Sequences were simulated to length 2600 base pairs (the approximate length of the HIV envelope gene) using the *env* mutational model from sim.coal.phybreak() in the R package biophybreak (Lundgren et al. 2022), which implements a GTR model with discrete Gamma rate heterogenity and empirically-determined base frequencies, transition rates, and proportion of invariant sites. Time-calibrated trees were inferred with MrBayes (Ronquist et al. 2012) using a strict clock, a GTR+Γ model, and priors on the substitution rate and root age that were consistent with the simulation values. An outgroup was added to each tree (sampled at the most recent ingroup sampling time, and attached 50% deeper than the ingroup MRCA) before sequence simulation and removed after timetree inference. We found that trees could be inferred with enough fidelity to lead to similar inferences of transmission time as on the ‘true’ coalescent trees (Fig. S11). This sequence simulation assessment does indicate that our coalescent trees are within a realm representing potential HIV data. It does not prove, however, that trees of such quality could be inferred from real-world HIV datasets, which almost certainly include much more heterogeneous processes of evolution and sampling. We return to this theme when discussing results from our empirical analyses.

### Empirical tests

To assess how well our Markov time-slice model works on real-world data, we used previously-established transmission pairs with some reported clinical information about transmission time and direction. The known case data allowed for comparisons against our inferences from only the sequence data. We began with the existing large compilation of HIV transmission pairs from Leitner and Romero-Severson (2018).

We then considered only pairs that met several criteria, intended to provide high-quality test cases for our method: previously-reported para/polyphyletic tree topology, at least ten tips on the tree (clonal sequences) for a given genomic region from each host, and tips from at least two different sampling times. From this subset of pairs, we selected three that had distinct transmission modes (or, risk factors), preferring pairs with more tips, more sampling times, and more precise clinical information about transmission direction and timing. The three case studies are summarized in Results, with more details in Table S1.

For each transmission pair, the alignment was downloaded from the “Special Interest Alignments” section of the LANL HIV database at https://www.hiv.lanl.gov/content/sequence/HIV/SI_alignments/datasets.html. We used BEAST v.1.10.4 (Suchard et al. 2018) to infer a posterior set of time-calibrated trees. This included a GTR+Γ model to allow variation in the substitution rate among sites (Yang 1993; Leitner et al. 1997), and an uncorrelated lognormal relaxed clock to allow substitution rate heterogeneity across the tree (Drummond et al. 2006). We chose a normally-distributed prior on the clock rate, with mean and standard deviation (0.008 and 0.004 subs/site/yr, respectively) computed from data on the evolutionary rates in the relevant genomic region (*env*, for these case studies) for eleven patients (Zanini et al. 2015). For the direction-naive trees, we used a constant population size coalescent prior on the node times. For the direction-aware trees, following the methods of Vrancken et al. (2014), we specified which sequences came from the source and which from the recipient, and we and provided a prior distribution on when transmission could have occurred. The node time prior was then based on a constant population size within the source, and a population growing logistically within the recipient beginning at the transmission time.

We removed the reference outgroup sequences in the original alignments before timetree construction. The outgroup to ingroup lineages must have passed through many different hosts, so the evolutionary rates along their long branches would likely be even more heterogeneous than in the two focal hosts. We confirmed, however, that the rooting produced by BEAST with tip dates and a clock model agreed with the outgroup-based rooting produced by RAxML (Stamatakis 2014), which we used to infer maximum likelihood trees with branch lengths in expected number of substitutions per site (Fig. S4).

Timetree inference also required a prior on the root age. The time the source was infected provides some guidance, but the root could be older if the source was infected with multiple surviving lineages, or the root could be younger if only one founding lineage left descendants in the eventual sample. We used an offset-lognormal root age prior to broadly encompass the source’s infection time, as reported in each case history. The mean, standard deviation, and offset of this distribution were: (4, 4, 0.600) years for the MTCT pair, (3, 4, 0.044) years for the MSM pair, and (6, 3, 0.052) years for the Knife pair. For each, the offset was determined by the earliest sampling date.

Each BEAST run continued for 10^8^ MCMC generations. Removing the first quarter as burn-in was more than sufficient, as judged by visual examination of the trace. An ESS of at least 400 was obtained for all key parameters in all cases, as reported by Tracer.

For each case study, we sampled 50 trees randomly from the posterior set and fit our Markov time-slice model to each. Model fitting proceeded as for the simulation tests, except that a smaller time slice (10 days instead of 1 month) was used for the MSM pair because of the short time elapsed between transmission and sampling. Results for the phylogical window and coalescent model are reported across the entire posterior set of trees.

## Acknowledgements

This work was supported by National Institutes of Health grant R01AI087520 to TL. We benefitted from R. Heil’s coding efforts on a related project, supported by a summer student fellowship from LANL’s Center for Nonlinear Studies. Members of the MolEvolEpid team in T-6 provided valuable feedback throughout this project. We are grateful to B. Vrancken for providing example BEAST code to infer a phylogeny along with a transmission event.

## Data Availability

HIV genetic sequence data are available from the “Special Interest Alignments” section of the LANL HIV database, at https://www.hiv.lanl.gov/content/sequence/HIV/SI_alignments/datasets.html. Code implementing the Markov time-slice model is available at https://github.com/MolEvolEpid/timeslice.

---

**Figure S1:**
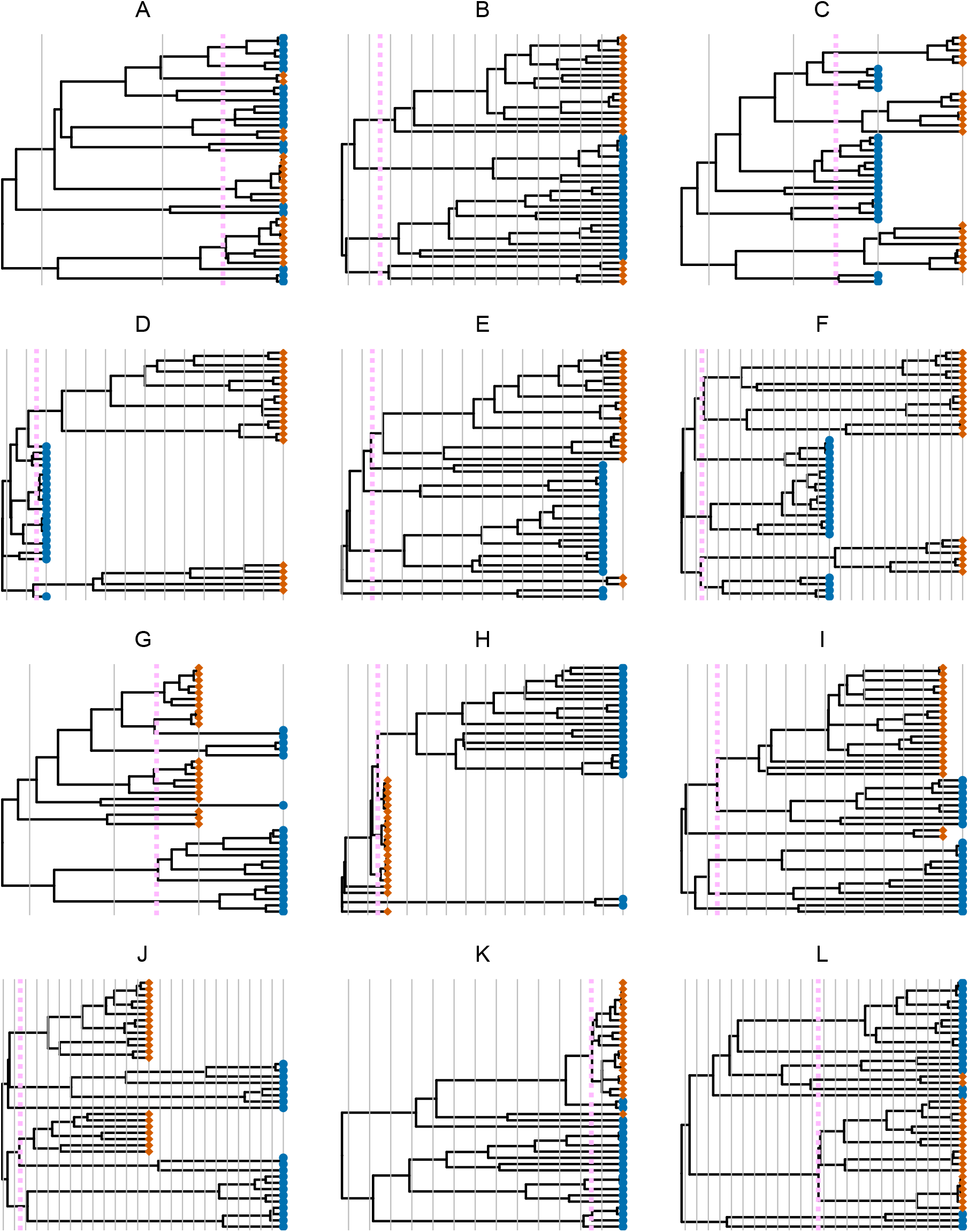
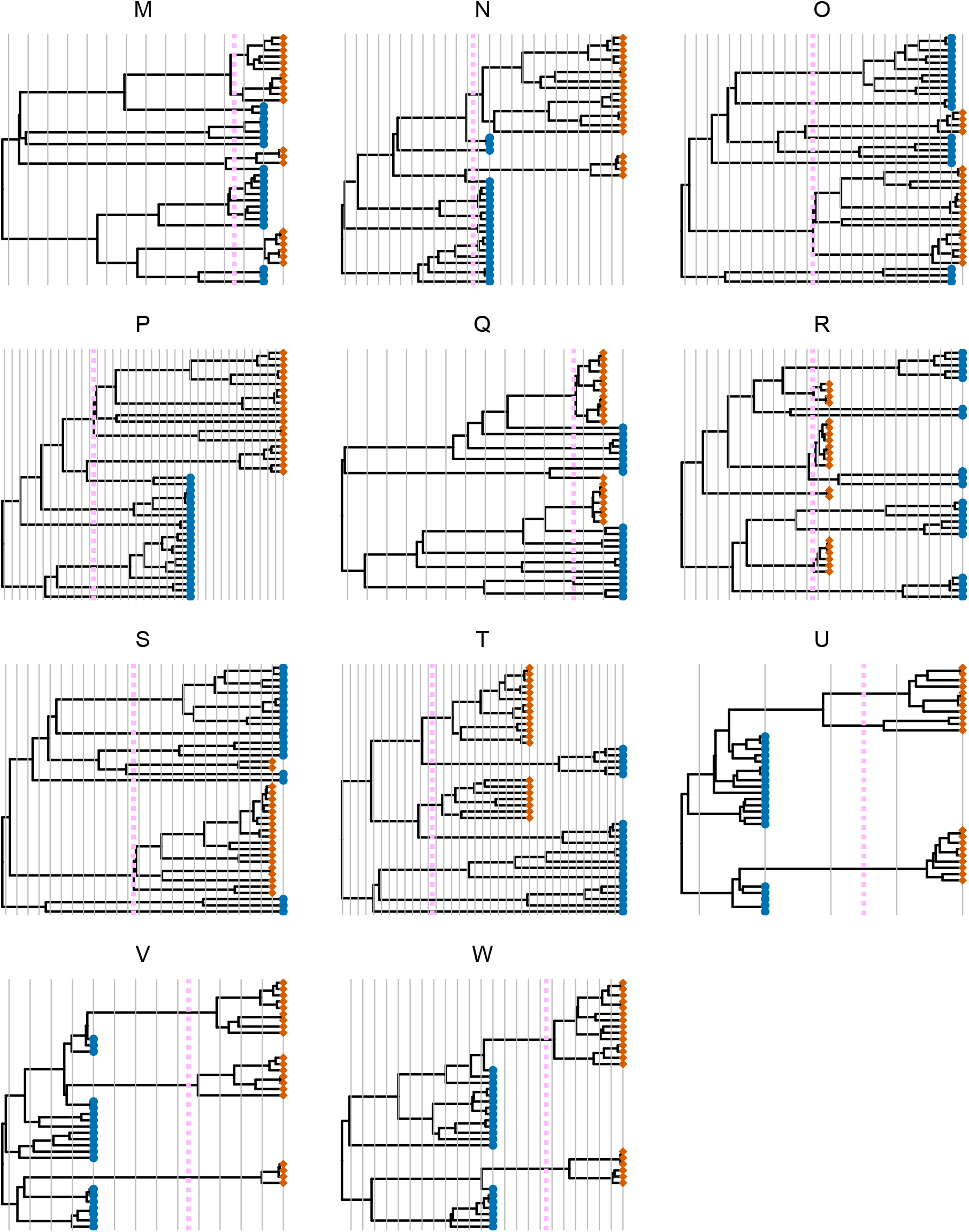
One representative PP tree from each simulated transmission history (Fig. 2). Tips from the source are blue circles, tips from the recipient are orange diamonds, and the transmission time is marked by the dashed pink line. Thin gray lines separate the one-month time slices used by the inference model.

**Figure S2:**
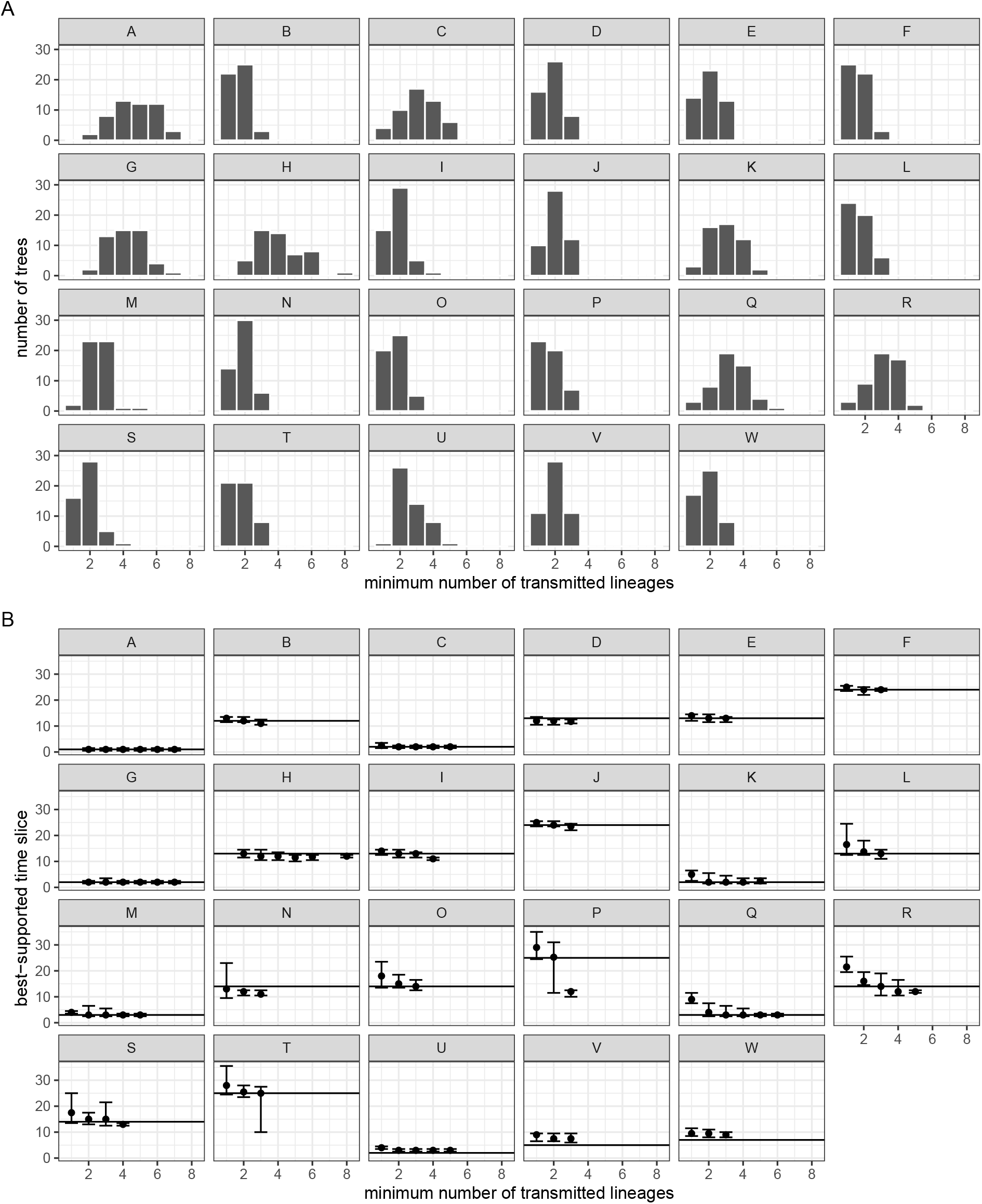
Minimum number of transmitted lineages, for each simulated history, counted as the number of distinct clades of recipient tips on a tree. (A) Bar plot across each set of 50 trees. (B) Accuracy of inferred transmission time, across trees with the indicated minimum number of transmitted lineages. The horizontal black line marks the true time slice, while points and whiskers show the median and 90% CI estimate, respectively.

**Figure S3:**
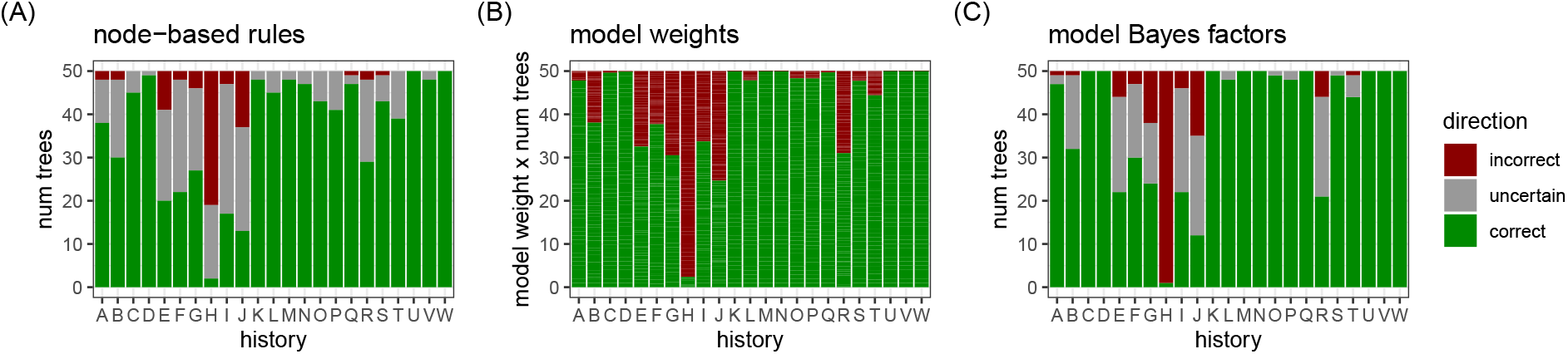
Inference of transmission direction on simulated data. For each simulated transmission history (Fig. 2A) with 50 replicate trees, colors indicate the number of trees for which the inference of transmission direction was correct, incorrect, or uncertain. (A) When node states are assigned according to a heuristic set of rules, the host state at the root is inferred to be the source (Fig. 1B). (B) In our time-slice model (Fig. 1D), the total weight for each transmission direction, across all time slices, is computed for each tree. These weights are then simply summed across trees. (C) Alternatively, the model weights for a tree can be used to compute a Bayes factor for the transmission direction. Compared with (B), panel (C) reports the transmission direction inference as uncertain for a tree if the Bayes factor is less than 3 (Kass and Raftery 1995).

**Figure S4:**
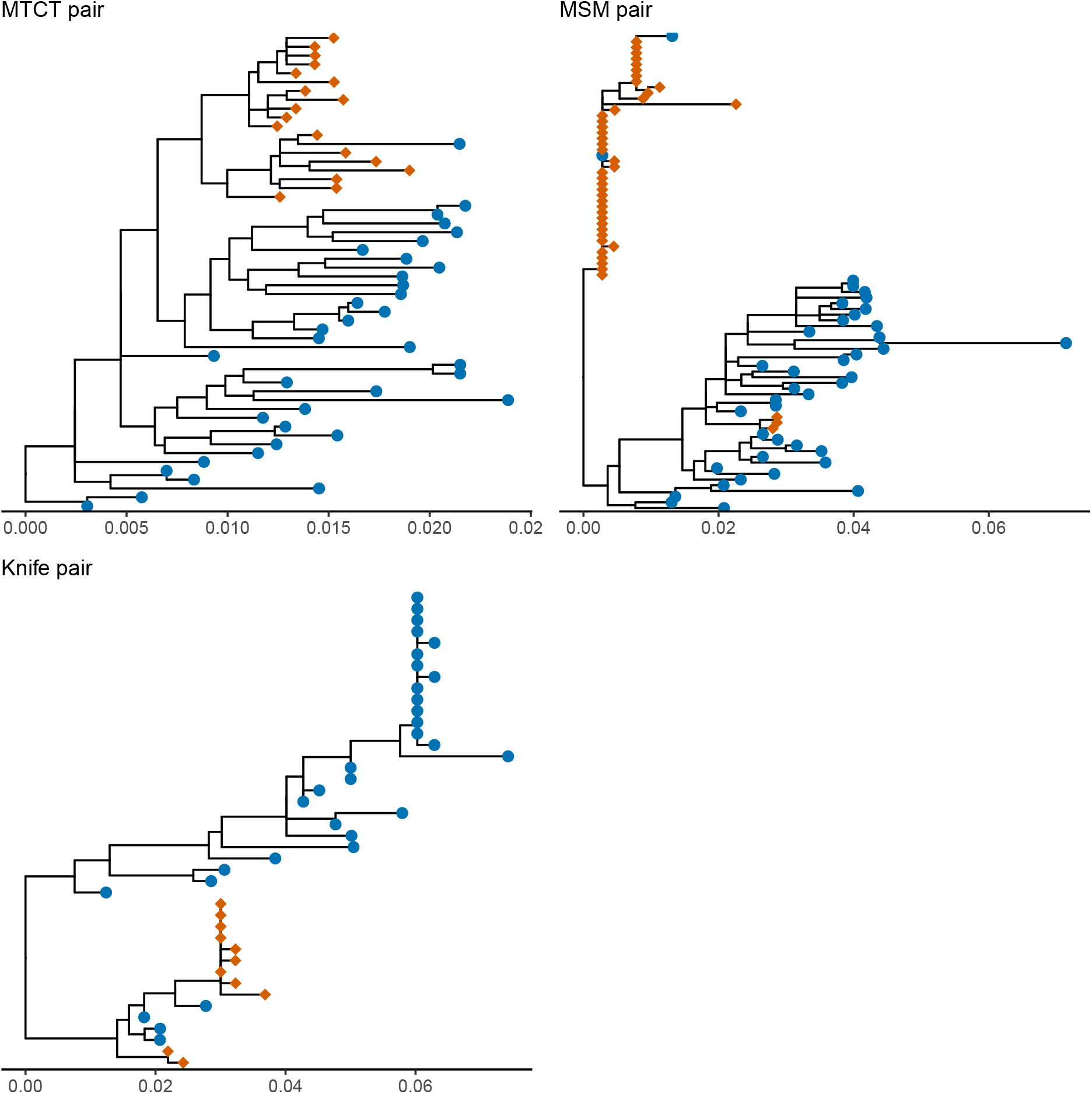
For the empirical case studies, maximum likelihood trees with branch lengths in units of substitutions per site. Each tree was inferred with RAxML (Stamatakis 2014). Outgroups were used for rooting and then removed for plotting. Sequences from the source are shown with blue circles, and from the recipient with orange diamonds.

**Figure S5:**
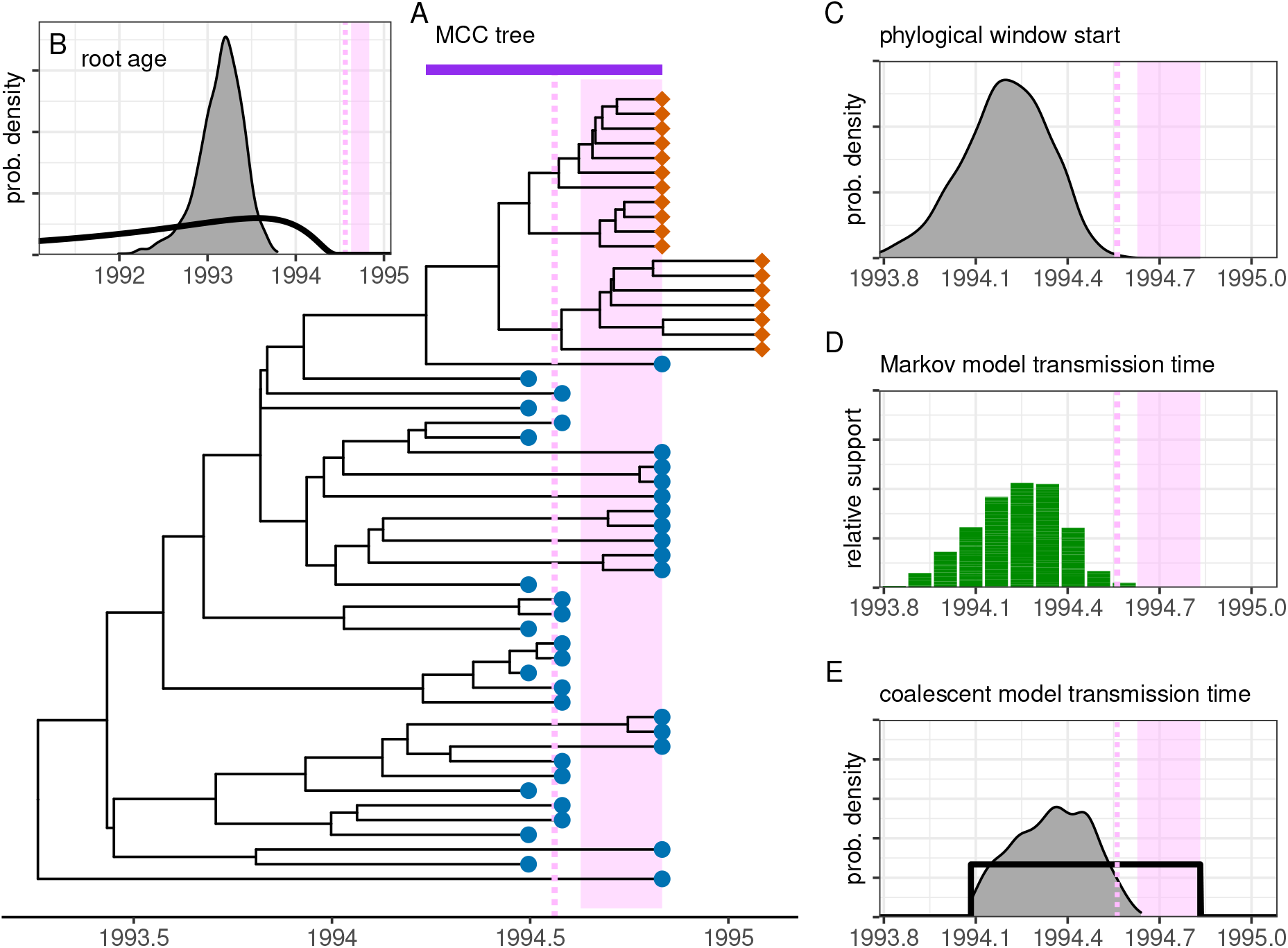
MTCT pair case study. These results are from a direction-informed phylogenetic inference, with an uninformative prior on transmission time that spans from the beginning of the pregnancy to the first HIV positive test (E). Components and vertical axis scales are as in Fig. 4.

**Figure S6:**
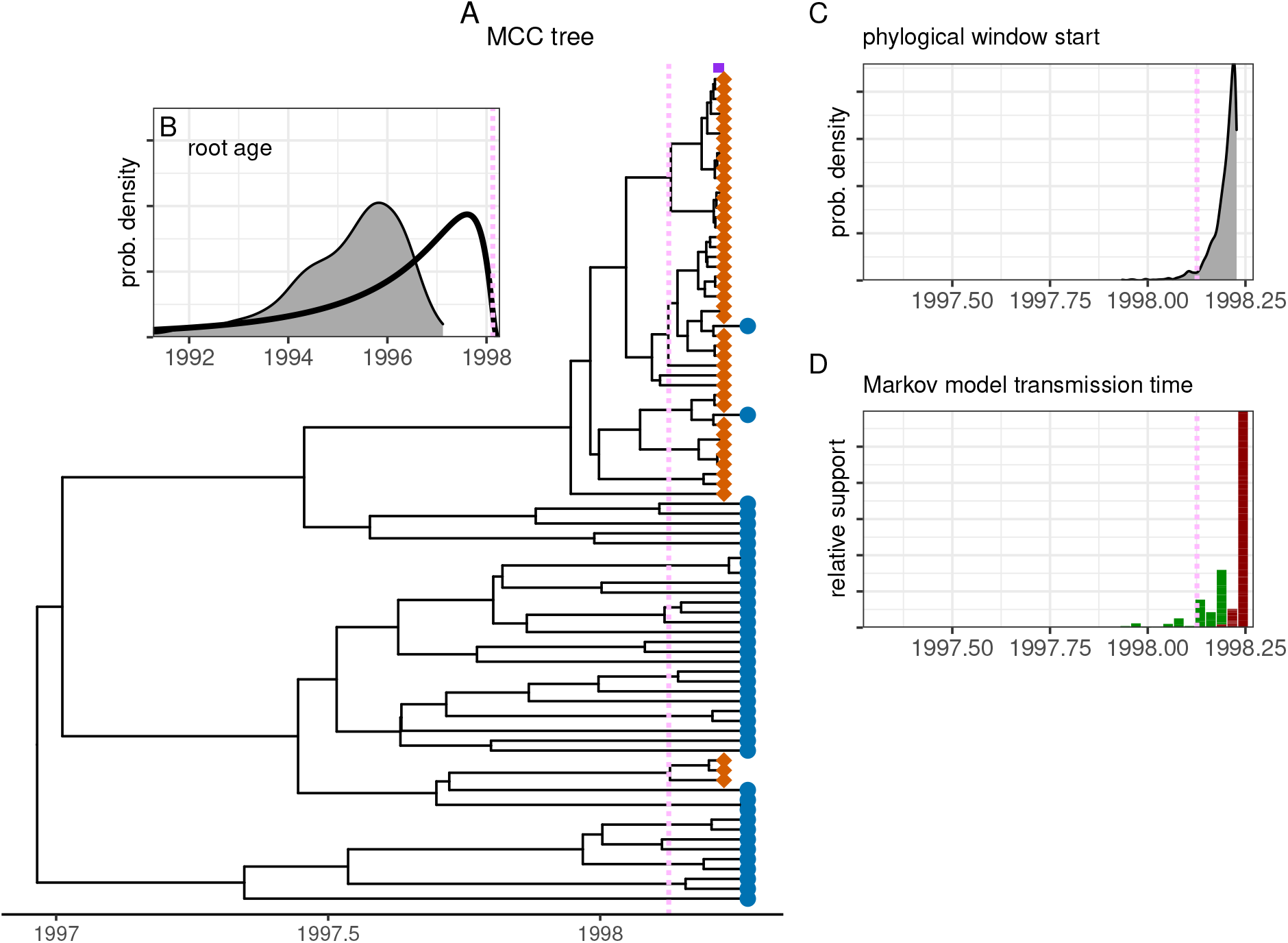
MSM pair case study. These results are from direction-naive phylogenetic inference. Vertical axis scales are the same as Fig. 5.

**Figure S7:**
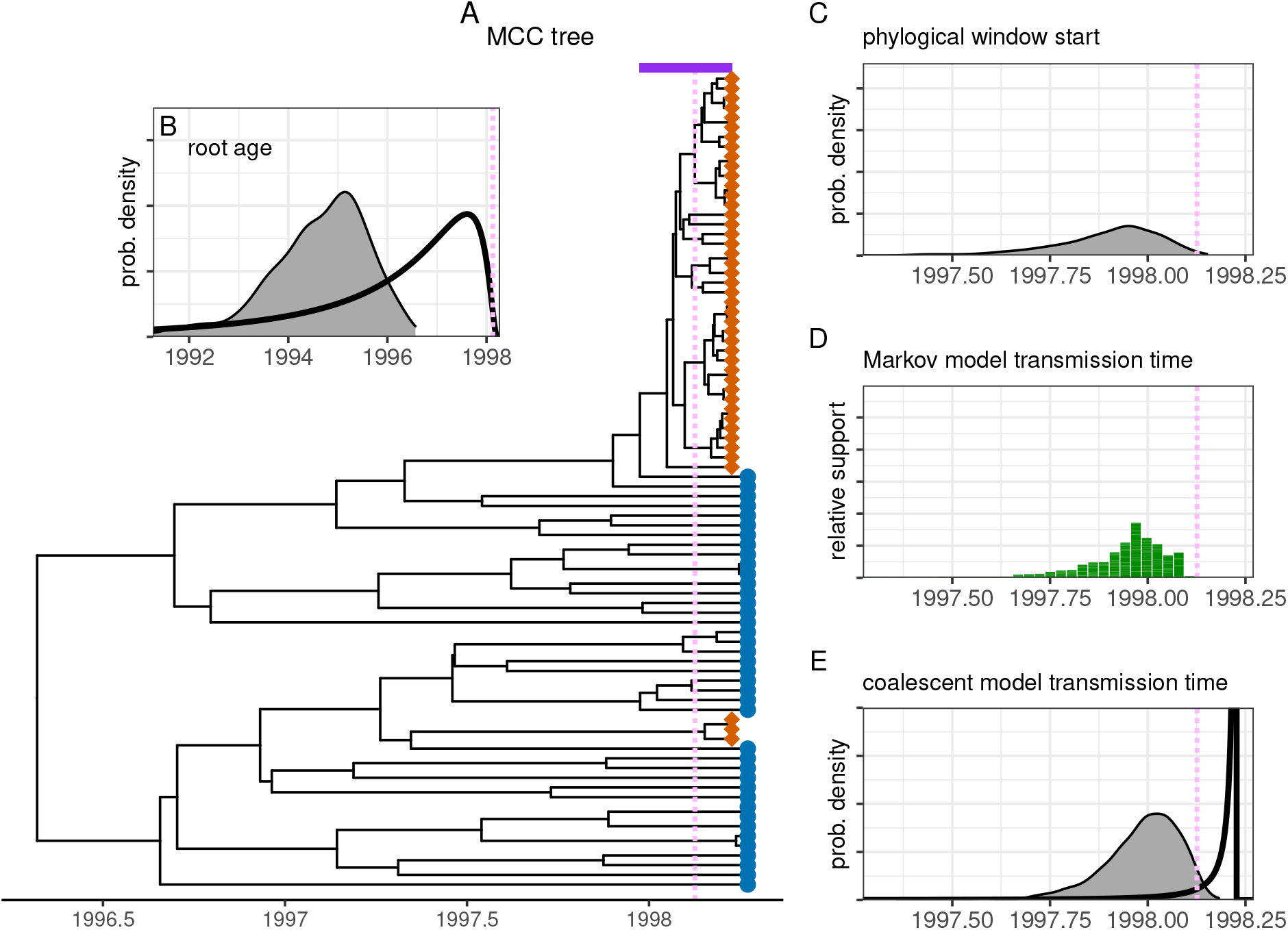
MSM pair case study. These results are from direction-aware phylogenetic inference, with an informative prior on the transmission time (E) based on time inferred by other means in the original study. Vertical axis scales are the same as Fig. 5.

**Figure S8:**
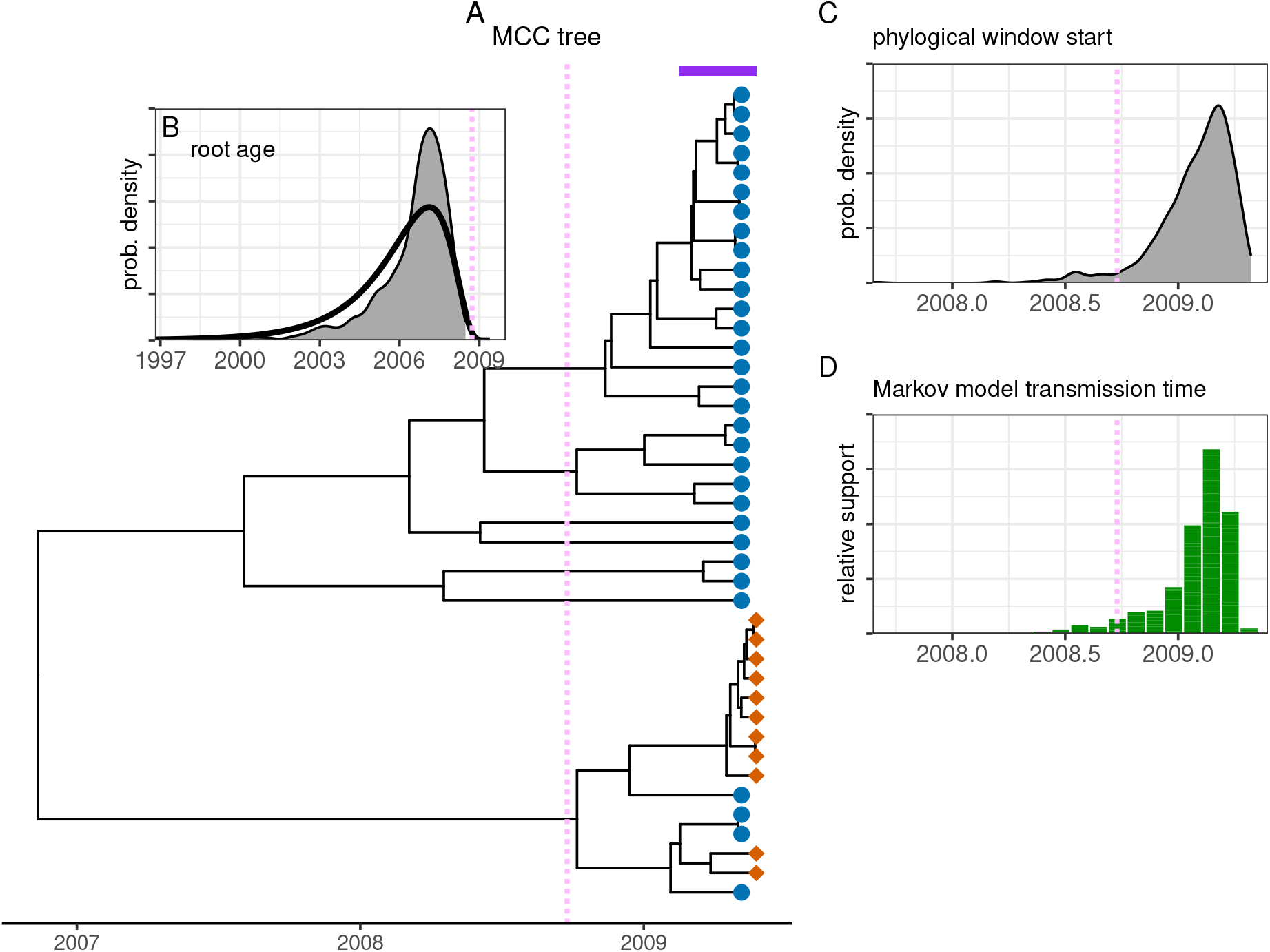
Knife pair case study. These results are from direction-naive phylogenetic inference. Vertical axis scales are the same as Fig. 6.

**Figure S9:**
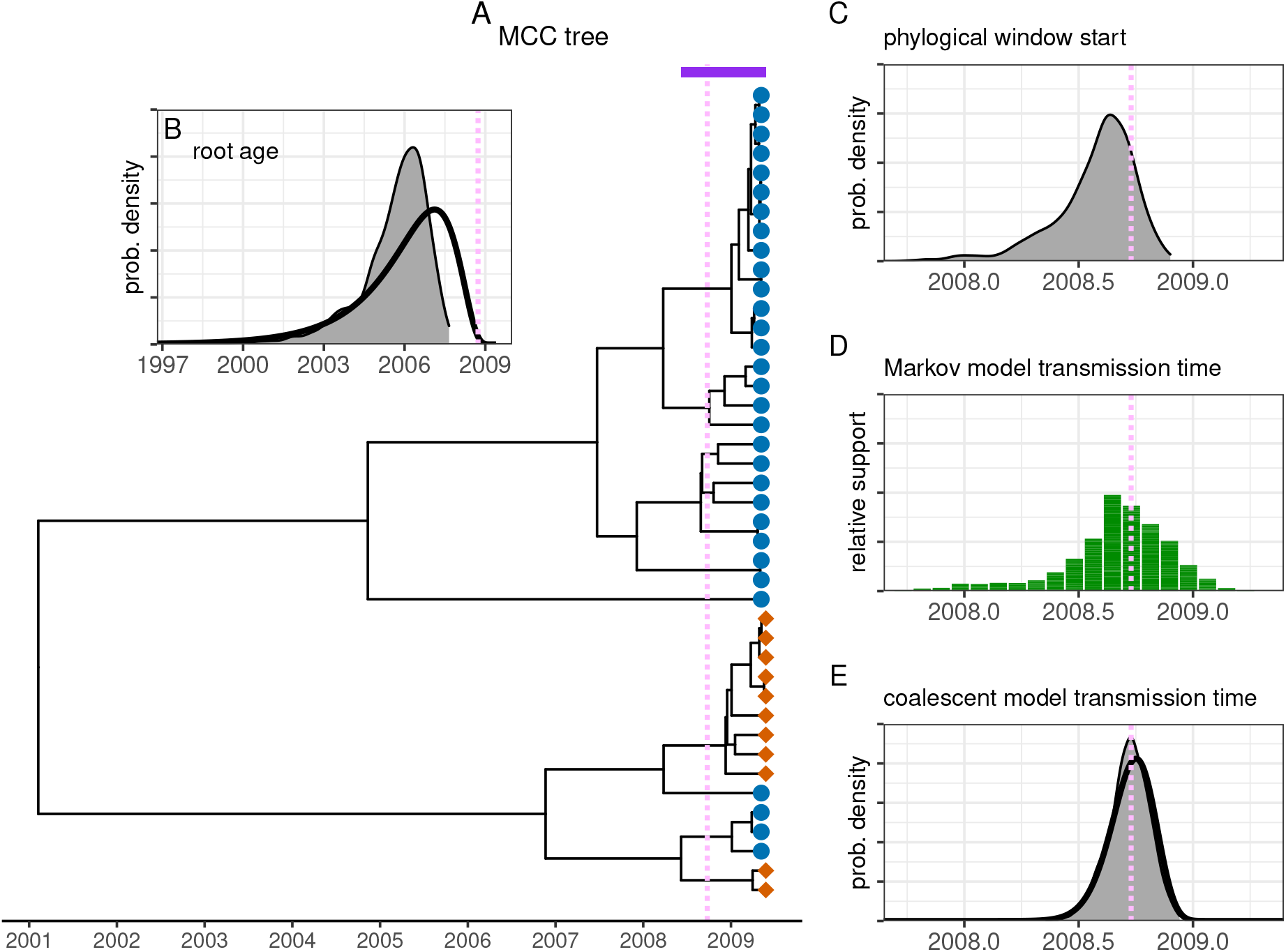
Knife pair case study. These results are from direction-aware phylogenetic inference, with an informative prior on the transmission time (E) based on the known date of transmission. Vertical axis scales are the same as Fig. 6.

**Figure S10:**
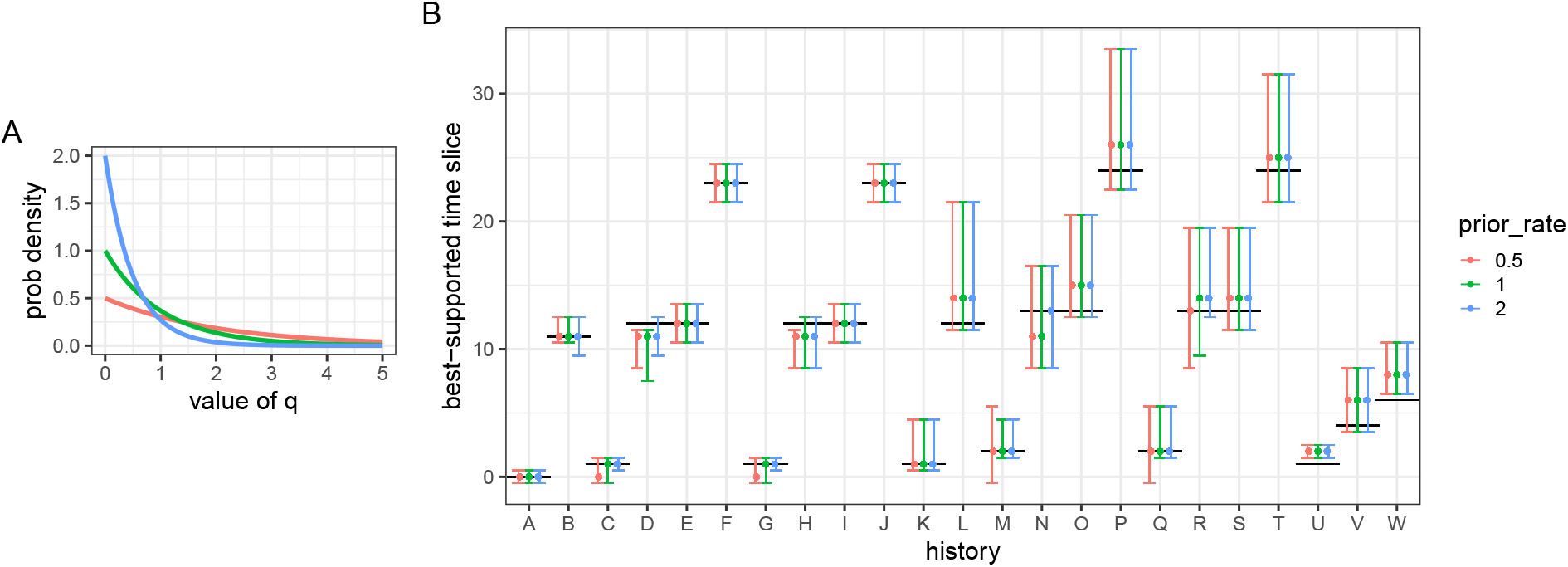
Effect of the prior on the time-slice model rate parameter *q*. Our main results all use an exponential prior on *q* with a rate of 1 per year (green). Results from either doubling (blue) or halving (red) the rate used in the exponential prior are also shown here. (A) Three possible prior distributions on *q*. (B) The best-supported time slice for each simulated history. Points show the median across trees, whiskers show the 90% CI, and black lines show the true value. Green results correspond to those shown in Fig. 3.

**Figure S11:**
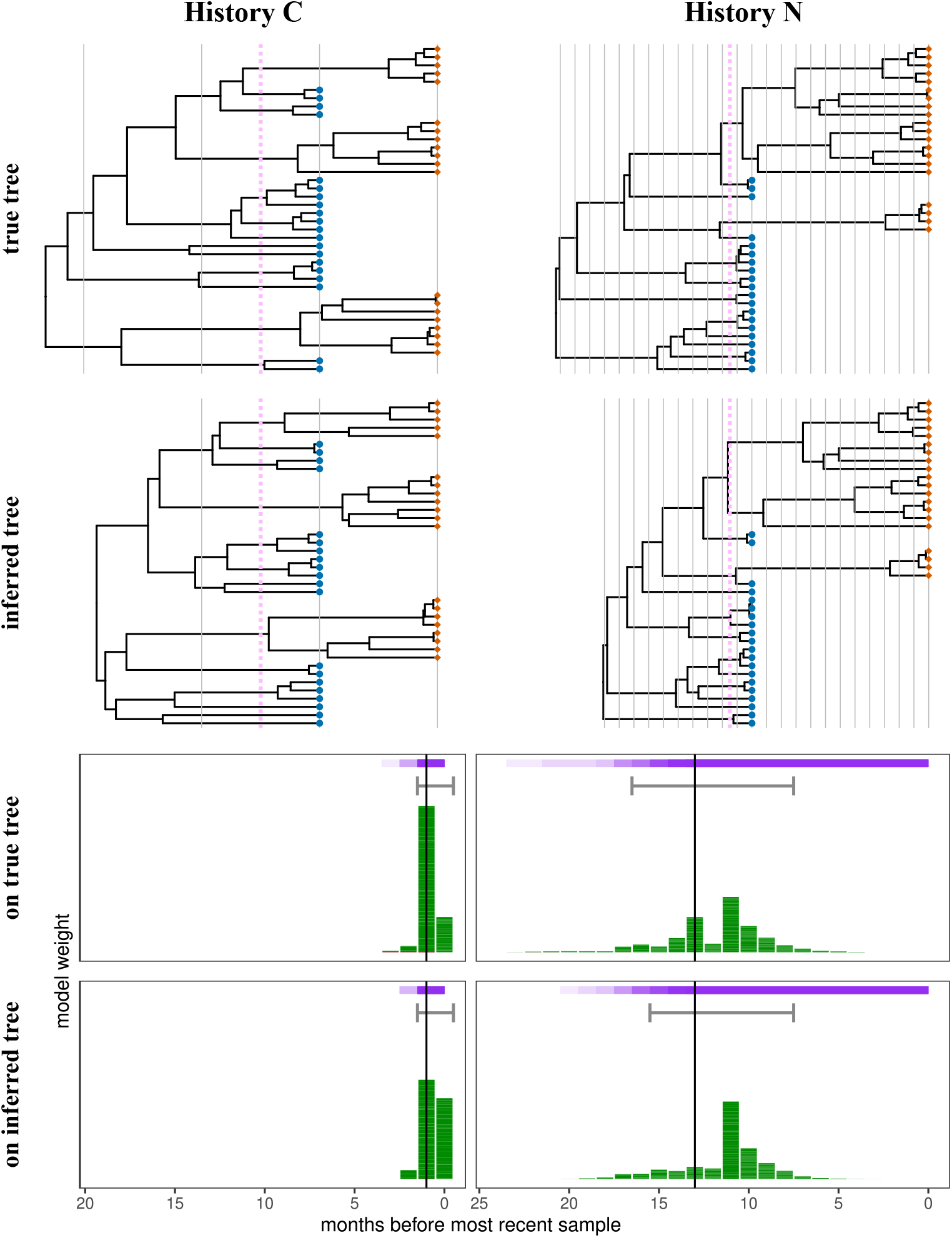
Tree and transmission time inference when sequences are simulated on coalescent trees. Results are shown for two histories, C and N (Fig. 2). First row: one example coalescent (‘true’) tree for each history. Second row: MCC tree inferred by MrBayes from an alignment simulated on that coalescent tree. Third row: transmission time inferred on the coalescent trees (also seen in Fig. 3, where figure elements are explained). Fourth row: transmission time inferred on the MCC trees for all coalescent trees.

**Table S1:** Case study details. Label is used in this paper. Citation is to the original study of the transmission cluster. Cluster ID is used in the HIV database at hiv.lanl.gov. Patient ID is for the source and recipient, which the LANL HIV database links to each NCBI accession. Genomic region provides coordinates and gene name of data in the alignment. Number of sequences is provided for the source and recipient.

| Label | Citation | Cluster ID | Patient ID | Genomic region | Num seqs |
| --- | --- | --- | --- | --- | --- |
| MTCT pair | Simonich et al. (2016) | MB_pair_520(1390) | 90186, 90187 | 6225–8795 (env) | 36, 18 |
| MSM pair | Frange et al. (2013) | SM_pair2(1040) | 60590, 60578 | 7005–7643 (env) | 40, 44 |
| Knife pair | Kao et al. (2011) | knife_fight(630) | 38458, 38457 | 7018–7502 (env) | 31, 11 |

